# The effects of cryptic diversity on diversification dynamics analyses in Crocodylia

**DOI:** 10.1101/2025.01.23.632920

**Authors:** Gustavo Darlim, Sebastian Höhna

## Abstract

Incomplete taxon sampling due to underestimation of present-day biodiversity biases diversification analysis by favouring slowdowns in speciation rates towards recent time. For instance, in diversification dynamics studies in Crocodylia, long-term low net-diversification rates and slowdowns in speciation rates have been suggested to characterise crocodylian evolution. However, crocodylian cryptic diversity has never been considered. Here, we explore the effects of incorporating cryptic diversity into a diversification dynamics analysis of extant crocodylians. We inferred a time-calibrated cryptic-species-level phylogeny using cytochrome *b* sequences of 45 lineages compared with the formally recognized 26 crocodylian species. Diversification rate estimates using the cryptic-species-level phylogeny show increasing speciation and net-diversification rates towards the present time, which is contrasting to previous findings. Cryptic diversity should be considered in future macroevolutionary analyses, however representation of cryptic extinct taxa represents a major challenge. Additionally, further investigation of crocodylian diversification dynamics under different underlying genomic data is encouraged upon advances in population genetics. Our case study adds to diversification dynamics knowledge of extant taxa and demonstrates that cryptic species and robust taxonomic assessment are essential to study recent biodiversity dynamics with broad implications for evolutionary biology and ecology.

## 1 Introduction

Macroevolutionary diversification dynamics analyses are fundamental for the understanding of evolutionary processes underlying historical biodiversity through time (1; 2; 3). From a reconstructed time-calibrated phylogeny, diversification dynamics analyses are able to track shifts of speciation and extinction rates. It additionally permits, for instance, to investigate possible intrinsic and/or extrinsic factors (or drivers) shaping the evolutionary history of a given group (4; 2; 3; 5). Therefore, taxon sampling —or lineage sampling— is intimately related to diversification rates estimation and diversification model selection (6; 7; 8). Incomplete taxon sampling biases diversification analyses by, for instance, favouring slowdowns in speciation rates towards recent-time (6; 9; 10). Incomplete taxon sampling is especially evidenced by the lack of representation of cryptic diversity in diversification studies due to inherent complexity of species concepts (11; 12; 13). Taxonomic assessment of cryptic species is contentious, however limiting species recognition under traditional taxonomic practices (i.e. morphology-based diagnosis) commonly makes cryptic diversity unavailable to different areas of biodiversity research (14). Ultimately, neglecting cryptic diversity underestimates present-day biodiversity and subsequently hampers a comprehensive understanding of the biodiversity science and its different areas including taxonomy, conservation, ecology, biogeography, and macroevolution (15; 14; 12; 16; 17; 18).

Crocodylia is a good example of a clade in which the representation of extant diversity is contentious subject to contrasting species recognition approaches (e.g. morphological or molecular-based) (25; 19). With an estimated age of approximately 95 million years (26; 27; 28; 29; 30; 31) and exceptional fossil record (32; 33; 34; 35), Crocodylia represents the crown group of Crocodyliformes, currently represented by semi-aquatic, ambush predators that inhabit freshwater, estuarine, and marine waters in tropical and subtropical regions of the globe (36). Crocodylia is composed by three main lineages, Alligatoridae, Crocodylidae, and Gavialidae (37), and with an extant diversity represented by 26 formally recognized species (23; 24; 38; 22). Preliminary evidence suggests the presence of two additional species (i.e. *Crocodylus halli* and *Osteolaemus afzelii*), although formal validation and description of those forms are still needed (25; 24; 38; 39). Low extant diversity and superficial morphological similarities with ancient forms have been used equivocally to refer to crocodylians as “living fossils” (40). This term places the understanding of crocodylians as evolutionary static forms or remnant populations of extinct taxa that once lived in the deep geological past (26; 25). However, such argument is contrasting to the diverse crocodylian fossil record (32; 35; 41; 42; 43). The term “living fossil” is furthermore problematic as it supports a conservative approach in crocodylian systematics concerning species recognition, which is in fact challenged by the discovery of cryptic species complexes (25). Multiple lineages of cryptic species complexes among crocodylians have been recognized due to advances in species delimitation methods in molecular studies, thus indicating a considerably higher extant diversity than otherwise suggested by morphology-only observations. Genetically distinct populations have been detected among the three main crocodylian lineages, such as in gavialids (44; 45), crocodylids (23; 24; 46; 38; 47; 22; 48; 49; 50), and alligatorids (51; 52; 53; 20; 19; 54; 55; 21) (figure 1). However, crocodylian cryptic diversity awaits further taxonomic assessment (25; 39).

**Figure 1:**
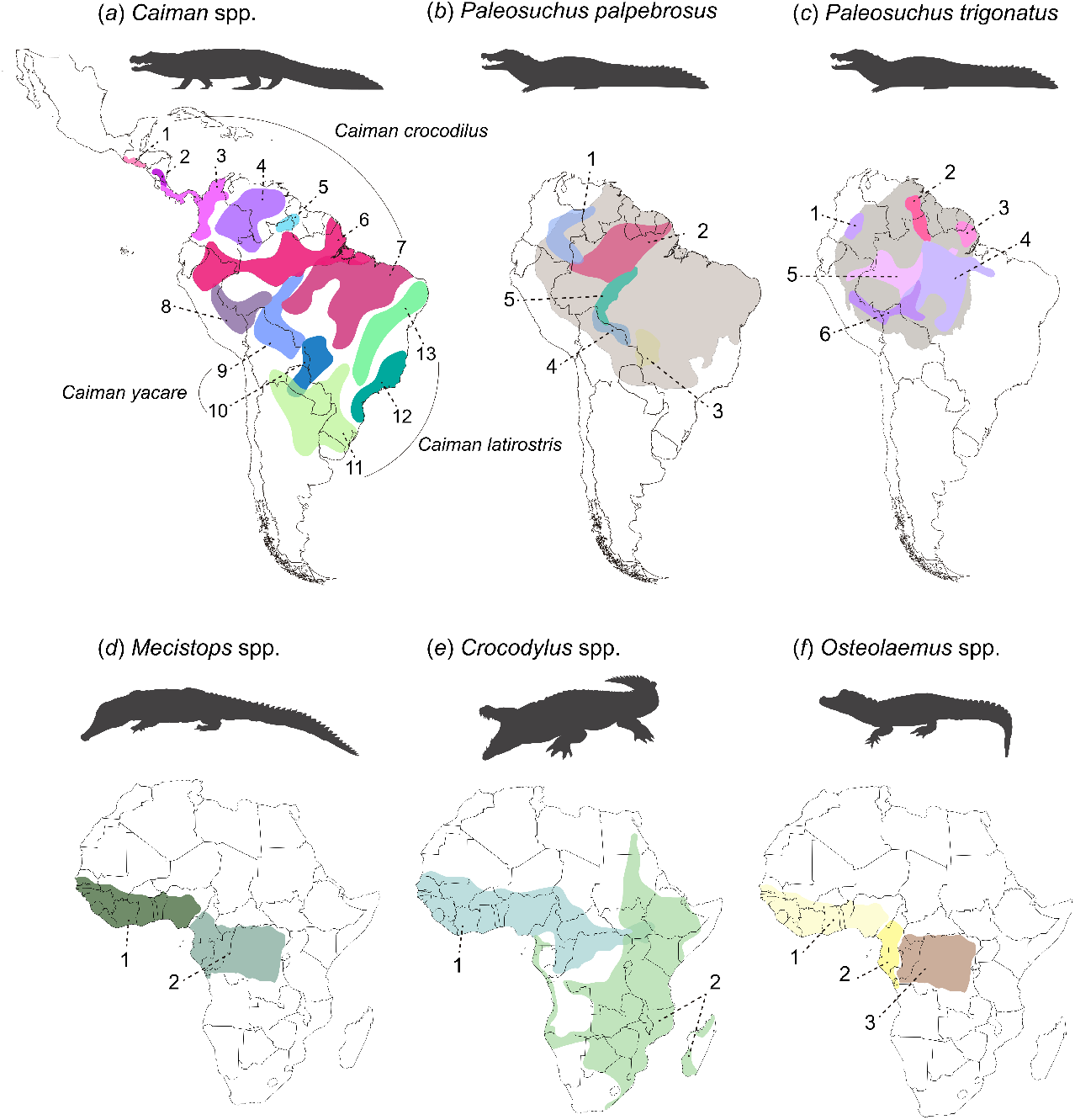
Maps showing the distribution of cryptic species in Crocodylia. Colored ranges in the map are numbered to indicate distribution of populations composing cryptic species complexes. (a) Distribution of *Caiman* species based on (19). *Caiman crocodilus* species complexes are indicated by numbers 1-8 (1, *C. c*.*chiapasius*; 2, *C. c*. cf. fuscus; 3, *C. c*.*fuscus*; 4, C. c. Orinoco; 5, *C. c*. Upper Branco; 6, *C. c*. Amazonia; 7, *C. c*. Brazilian shield; 8,*C. c*. South West Amazon); *Caiman yacare* species complexes are indicated by numbers 9-10 (9, *C. yacare* Bolivia Madeira; 10, *C. yacare* Pantanal); *Caiman latirostris* species complexes are indicated by numbers 11-13 (11, *C. latirostris* Paraná; 12, *C. latirostris* Doce; 13, *C. latirostris* São Francisco); (b) Distribution of *Paleosuchus palpebrosus* cryptic species (1, *P. p*. Orinoco; 2, *P. p*. Amazonas; 3, *P. p*. Pantanal; 4,*P. p*. Bolivia; 5,*P. p*. Madeira), based on (20; 21). Shaded gray area indicates distribution delimitation of *P. palpebrosus* based on IUCN; (c) Distribution of *Paleosuchus trigonatus* cryptic species (1, *P. t*. West Orinoco; 2, *P. t*. Upper Branco; 3, *P. t*. Coastal; 4, *P. t*. East Amazon; 5, *P. t*. West Amazon; 6, *P. t*. South West Amazon), based on (20; 21); Shaded gray area indicates distribution delimitation of *P. trigonatus* based on IUCN; (d) Distribution of *Mecistops cataphractus* (1), and *Mecistops leptorhynchus* (2) based on (22); (e) Distribution of *Crocodylus suchus* (1), and *Crocodylus niloticus* (2) based on (23); and (f) Distribution of *Osteolaemus cf. tetraspis* (1), *Osteolaemus tetraspis* (2), and *Osteolaemus osborni* (3) based on (24). Maps and silhouettes are not in scale. Silhouettes were sourced from phylopic.org: *Caiman* and *Paleosuchus* (Armin Reindl) CC BY-NC 3.0 DEED license, *Crocodylus* (Steven Traver) CC0 1.0 DEED Public Domain license; and modified from photographs in Wikimedia Commons: *Mecistops cataphractus* (Leyo) reflected horizontally, CC BY-SA 3.0 CH DEED license, *Osteolaemus tetraspis* (Adrian Pingstone) Public Domain license.

The discovery of cryptic diversity illustrates that the recognition of extant crocodylian diversity based solely on traditional taxonomic practices (i.e. species-level definition) affects different areas of biodiversity research, including for example, conservation, systematics and macroevolution (24; 19; 54). Therefore, addressing evolutionary processes to discuss crocodylian extant diversity has been advocated among crocodylian systematists and conservation communities (24; 19; 54). Nonetheless, in the assessment status carried out by The International Union for Conservation of Nature (IUCN) Red List of Threatened Species, a Last Concern status is currently assigned to most of cryptic crocodylian lineages as they compose species complexes of widely distributed and abundant taxa (i.e. *Caiman crocodilus*). This is problematic as most of the detected cryptic species are in fact threatened by habitat modifications an warrant urgent conservation strategies and implementation of managing programmes (19; 55). At least seven extant crocodylians species are categorized as critically endangered and three species are categorized as vulnerable (39). Whereas most recent recognized species —*Crocodylus suchus, Mecistops leptorhynchus*, and *Osteolaemus osborni* — still awaits for further assessment on their conservation status (39), cryptic species detected among South American alligatorids (i.e. *Caiman* spp. and *Paleosuchus* spp.) have still not being addressed specifically. Therefore, it illustrates biases and limitations of sole use of traditional taxonomic practices (species-level) into capturing crocodylian extant diversity.

Similarly, a more comprehensive sampling of crocodylian extant diversity (i.e. including cryptic diversity) has not yet been addressed in other areas of crocodylian research, such as macroevolutionary studies. The most comprehensive sampling of extant crocodylian species in diversification dynamics analyses includes 26 extant species (56) (i.e. estimated molecular phylogeny used for the diversification dynamics analysis), whereas 23 species are included in Payne *et al*. (30) (i.e. metatree approach), and finally 14 species are included in the studies of Magee and Höhna (5) (i.e. based on a published phylogeny), and Solórzano *et al*. (57) (i.e. based on species occurrence). Except for Colston *et al*. (56), the aforementioned studies additionally use data from the fossil record. These diversification dynamic studies have detected long-term low net-diversification rates in Crocodylia (56; 57; 30; 5) with peaks of higher diversity observed in the late Paleocene, late Eocene, and Early Miocene (57). Following the Miocene, a sharp decline in speciation and increase of extinction rates leading to the current-day observed diversity have been recovered (30), and furthermore suggested that living crocodylians are overall in decline (5). Speciation rates of crocodylians have been furthermore characterized by slowdowns subsequent to a Miocene diversity peak, thus leading to the current observed diversity (5; 56; 30).

Therefore, in order to accommodate a more comprehensive sampling of crocodylian extant diversity, in the present study we used the most up to date available molecular data concerning well-established cryptic species to investigate diversification dynamics through time. We estimated and compared speciation, extinction and net-diversification rates between time-calibrated phylogenies including and excluding cryptic diversity using state-of-the-art diversification rate approaches (58) in `RevBayes` (59). Our study demonstrates that addressing cryptic diversity in diversification dynamics analyses affects speciation rates towards the recent time by mainly capturing lineages splitting events during the Pliocene and Pleistocene, differing from previous diversification dynamics studies. Additionally, our results show that discussions on crocodylian diversity detached from taxonomic definitions –thus not restricted to the species-level– allows for a more comprehensive understanding of crocodylian diversification dynamics. It is worth noting that the increase of speciation events recovered in our results does not equal less concern regarding conservation efforts towards crocodylian species. Instead, our results are complementary to discussions regarding the importance of recognizing cryptic diversity – and considering evolutionary processes into taxonomic acts, as advocated by previous studies (i.e. (19))– to appropriately address crocodylian extant diversity and to support more effective conservation strategies towards smaller and geographically restricted populations.

## 2 Methods

### 2.1 Molecular data

We collected molecular sequence data using GenBank accession numbers of the mitochondrial gene cytochrome *b* (*Cytb*) for 45 genetically distinct lineages. We followed recent studies on crocodylian population genetics in order to select the cryptic species for our analysis, in which we used one specimen per genetic cluster (haplogroup). Nomenclature for the different clusters follows (52; 20; 21) for *Paleosuchus* spp., and (51; 19) for *Caiman* spp. We provide a list of all species, accession numbers for the selected specimens, gene length (bp) and source studies for our dataset in the electronic supplementary material (Table S1). We used the software MUSCLE (Multiple Sequence Comparison by Log-Expectation (60)) for aligning the sequences using the software MEGA X (61).

### 2.2 Time calibrated phylogenetic inference and diversification dynamics analyses

We performed a joint analysis that simultaneously infers phylogenetic relationships, divergence age estimates, and rates of diversification through time. Therefore, the joint analytical approach avoids the selection of a single tree for a subsequent diversification dynamics analysis. Our species-level phylogeny corresponds to the phylogeny inferred by Darlim and Höhna (31). To obtain a comparable cryptic-specieslevel phylogeny, we followed the same protocol and used the same phylogenetic model and software. In detail, we specified the following statistical model. We partitioned the single gene into three data subsets according to codon positions. Then, we assumed that each data subset evolved under a general time reversible substitution model with gamma distributed rate variation across sites with four discrete rate categories (GTR+Γ) (62; 63). Substitution model parameters were independent for each data subset. We additionally applied partition rate multipliers drawn from a flat Dirichlet distribution. We applied two different sets of fossil node-calibrations for the following six nodes: Crocodylia, Alligatoridae, crownAlligatorinae, crown-Caimaninae, Longirostres, and Osteolaeminae. Calibration set 1 follows strictly recent recommended fossil calibrations (e.g. (64)) and the best practices protocol (65). Conversely, for comparison purposes, calibration set 2 consists of an alternative set fossil taxa that have been at some extent influencing the age of crown lineages and that are currently debated in crocodylian literature. Stratigraphic age intervals, justification for the selected fossils, parameters for the time constraints, and details of the calibration sets are provided in the electronic supplementary material (Tables S1 & S2) and in Darlim and Hohna (31). We used a relaxed-clock mixture model between an exponential, gamma and lognormal distribution to allow for substitution rate variation among lineages (31).

We estimated diversification rates (speciation and extinction rates) using the piecewise-constant birthdeath process (2; 3; 58). We divided time between 66 mya and the present into 100 equal-sized intervals. For each interval, we estimated separate speciation and extinction rates. We assumed a Horseshoe Markov Random Field smoothing prior for the rates between intervals (58). We emphasize that our joint analysis of diversification rates, divergence times, and phylogeny marginalizes over the phylogeny and divergence times when the diversification rates are the focal parameter and thus avoids to rely on a single estimated phylogeny.

We ran four replicate Markov chain Monte Carlo analyses for 500,000 iterations each and checked for convergence using the R package `convenience` (66). The phylogenetic analysis was performed in `RevBayes` (59). The results were summarized and plotted using the R package `RevGadgets` (67).

## 3 Results

Our estimated cryptic-species-level phylogeny is topologically congruent with the species-level phylogeny of Darlim and Hohna (31) (figure 2), and overall similar to those of previous molecular studies (e.g. (26; 27; 19)) (figures S2,S3,S6,S7). Divergence time estimates for both species-level and cryptic-species-level using the calibration set 2 retrieved significantly older ages that are in conflict with unambiguous crown fossil taxa (for details, refer to electronic supplementary material and Tables S3,S4). Conversely, age estimates when using calibration set 1 are overall similar to those of previous studies using well-justified calibrations (e.g. (26; 27)), thus more consistent with the evidence of the fossil record. Additionally, divergence age estimates for the cryptic-species-level phylogeny using the calibration set 1 closely match the age estimates of the species-level phylogeny with 95% credible intervals largely overlapping (Table S3). Interestingly, the overall pattern of diversification rates are the same retrieved for analyses using calibration set 1 (figures 3, S4) or calibration set 2 (figure S8). Here we discuss the results of our analyses based on the age estimates of the time calibrated phylogenies using the calibration set 1. A complete report of the results of our phylogenetic analyses, divergence age estimates, and diversification dynamics analyses are available in the electronic supplementary material.

**Figure 2:**
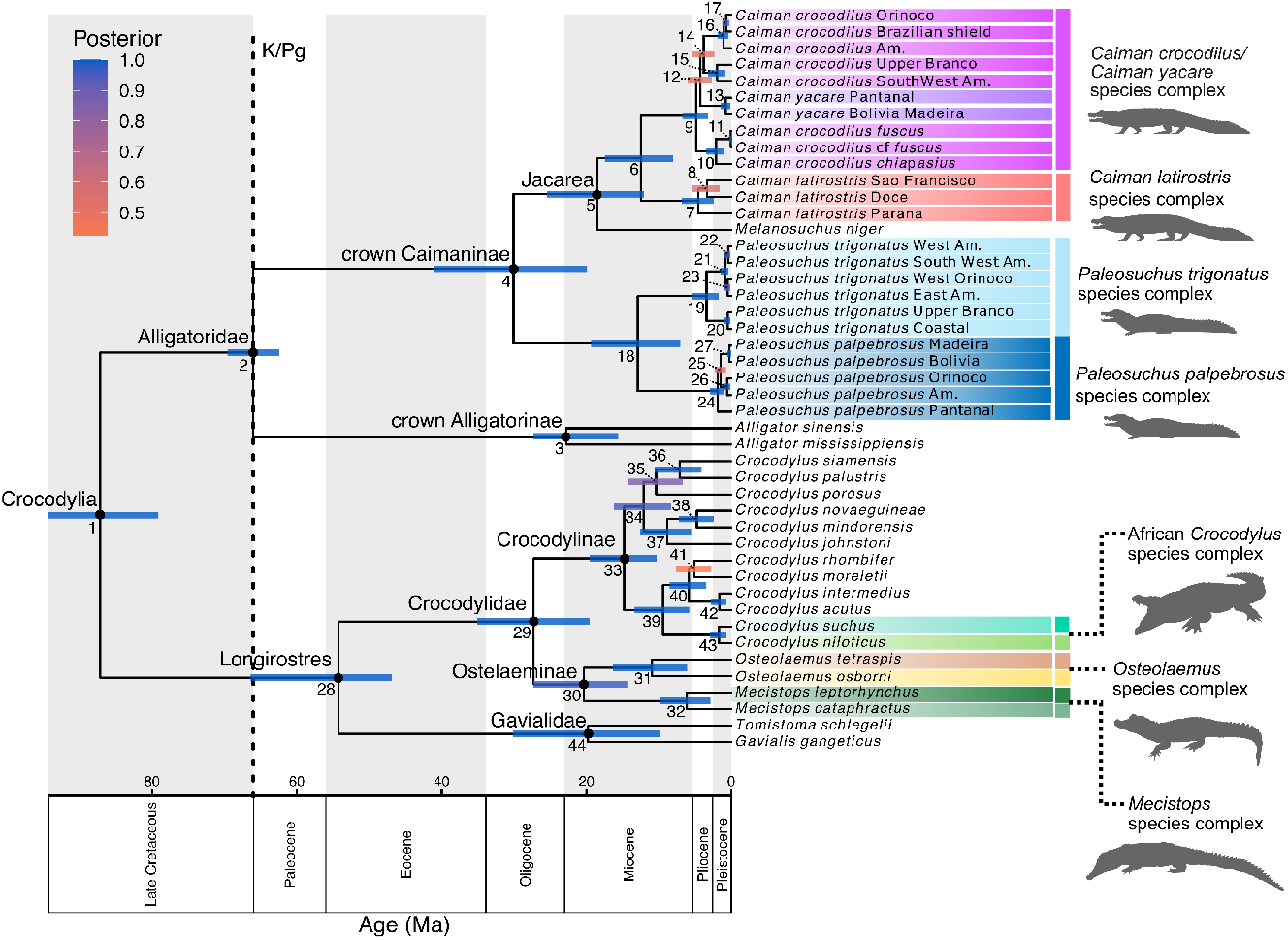
Time-calibrated cryptic-species-level phylogeny of Crocodylia using calibration set 1. Node bars represents 95% highest posterior density (HPD) intervals for age estimates. Color of node bars indicates posterior probability support following the index on the top left. Nodes are numbered, and their 95% highest posterior density intervals (HPD) can be be found at Table S3 in the electronic supplementary material. The phylogeny was plotted using the R package `RevGadgets` (67). Silhouettes sources are the same as specified in figure 1.

**Figure 3:**
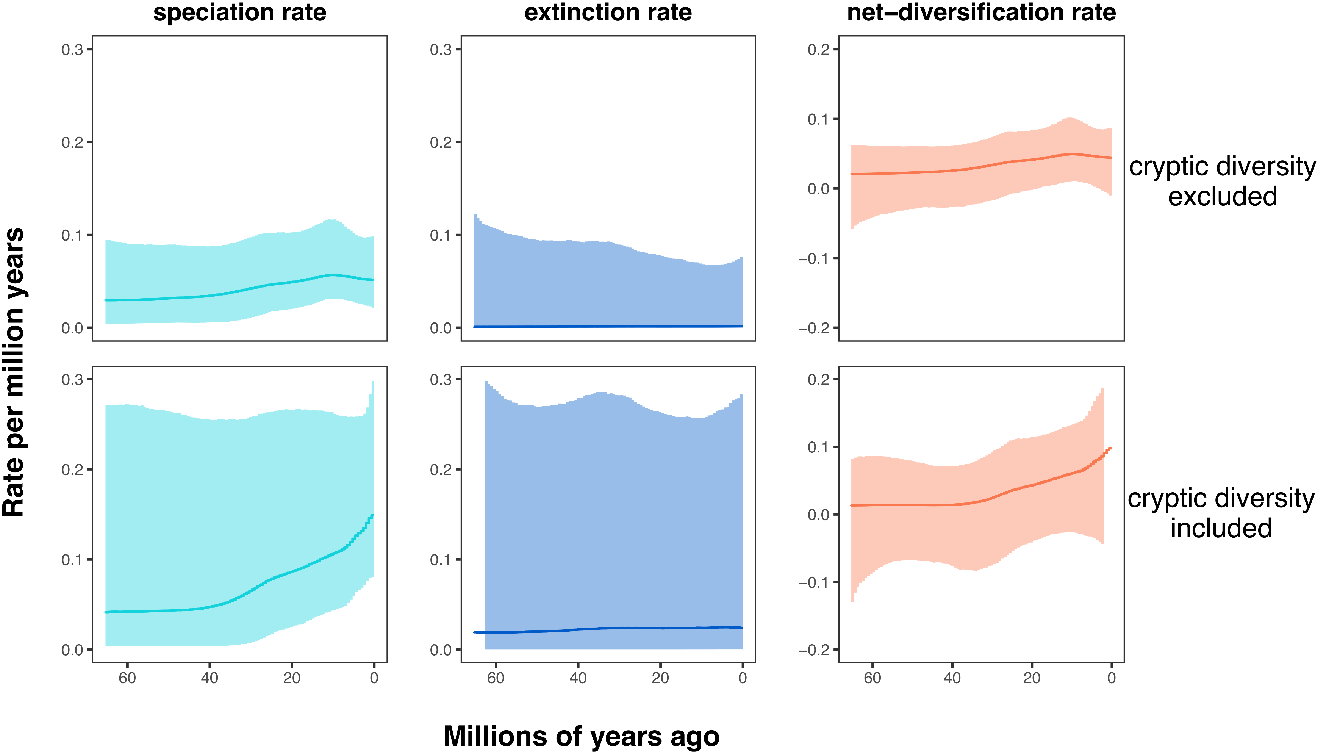
Diversification dynamics analyses in Crocodylia showing estimated speciation, extinction, and net-diversification rates across time based on data sets excluding (top row) and including (bottom row) cryptic diversity under calibration set 1. Analyses were performed under the piecewise-constant birth-death process with 100 equal-sized intervals between 66 million years ago and the present implemented in `RevBayes` (59). Results are summarized and plotted with the R package `RevGadgets` (67). Posterior probability of a speciation rate decrease for the specieslevel analysis between the peak at ∼ 10 mya and the present was *P* (*λ*(*t* = 10) *> λ*(*t* = 0)) = 0.61, which gives a Bayes factor of 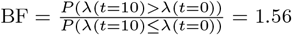,signifying weak support. Posterior probability of a speciation rate increase for the cryptic-species-level analysis between ∼ 40 mya and the present was *P* (*λ*(*t* = 40) *< λ*(*t* = 0)) = 0.89, which gives a Bayes factor of 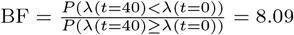,signifying substantial support. Solid line line represents the median.

The diversification dynamics analysis including the cryptic diversity in Crocodylia retrieved an increase in speciation and net-diversification rates compared to the analysis excluding cryptic diversity (figure 3). In the latter, curves of speciation and net-diversification rates through-time are overall constant (at approximately 0.02 − 0.05 speciation events per million years), with the exception of a slight peak at approximately 10 million years ago. Following this period, speciation and net-diversification rates slightly decrease towards the recent time (posterior probability of decrease of 0.61, Bayes factor of 1.56). Conversely, in the diversification analysis including cryptic diversity, curves of speciation and net-diversification rates continuously increase from approximately 40 million years to the present time, starting at approximately 0.05 at 40 million years ago to 0.15 speciation events per million years at the present (posterior median estimates, posterior probability of increase of 0.89, Bayes factor of 8.09, figure 3). Extinction rates are overall constant through time in both analyses and virtually zero for the diversification dynamic analysis excluding cryptic diversity.

## 4 Discussion

### 4.1 Cryptic diversity and crocodylian macroevolution

Results of our diversification analysis when using the time-calibrated extant phylogeny of Darlim and Höhna (31) (i.e. species-level phylogeny) show an overall low net-diversification rate, with a slight peak in speciation and net-diversification rates in the Miocene at approximately 10 mya, followed by a decrease on the speciation and net-diversification rates (figure 3), thus consistent with previous diversification studies (i.e. (56; 5)). In both of our diversification dynamics analyses (i.e. including and excluding cryptic diversity), a Miocene peak of speciation and net-diversification rates is consistent with the origination of modern crocodylian lineages at ca. 5 − 15 mya as also retrieved by previous molecular estimates (26; 24). Conversely, rates of speciation increases towards the recent time in our analysis when using the cryptic-species-level phylogeny, despite the extinction of many crocodylian lineages after the Miocene (e.g. (68; 57)). Our recovered increase in speciation and net-diversification towards the recent time (figure 3) is justified by the origination of sampled cryptic caimanine lineages during Pliocene and Pleistocene, combined with the observed diversification of crocodylids from the Late Miocene to the Pleistocene. Previous studies have suggested that the estimated Miocene time interval for the most recent common ancestor of modern lineages represents a suitable timeline for crocodylian speciation (24). Factors driving the extinction of many crocodylian species in the Late Miocene as evidenced in the fossil record (68; 32; 34), could have also contributed to the diversification of remnant crocodylian lineages via, for example, allopatric speciation (24). This is especially relevant with respect to the wide geographic distribution of extant *Caiman* and *Paleosuchus* species and the recent discovery of genetic diversity among different populations (19; 55; 20).

However, discussion regarding crocodylian speciation needs to be carefully evaluated. Diversification dynamic studies have demonstrated a complex interaction between biotic and abiotic factors underlying diversification dynamics in Crocodylia (57; 56; 30). Crocodylian speciation rates were suggested to represent diversity-dependent processes (57; 30), and potentially to be positively correlated to temperature (30), considering the increase of habitable areas for crocodylians due to the extension of warm latitudinal climatic belt (57). Extinction rates have been suggested to be negatively correlated to body mass disparity (57). Therefore, further investigation of those drivers of diversification in light of the impact of cryptic diversity of speciation and net-diversification rates retrieved by our study is encouraged, specially in combination with the fossil record. As recommended by previous studies (30; 5), the inclusion of fossils is key for understanding drivers of diversification, and detecting signals of extinction events. This matter is additionally evidenced by ongoing discussions on diversification dynamics analyses that are drawn solely from molecular phylogenies or in combination with the fossil record (e.g. (69; 70; 71; 10)). Diversification dynamics based solely on extant-only phylogenies are consistent with infinitely many diversification rate curves (non-identifiability (72)), however, only if completely arbitrary rate functions are allowed (73), and inferences of rapid changes are robust despite this non-identifiability (74; 75). Note that the specific model used here, the piecewise-constant birth-death process, has been shown to be identifiable (76; 77). The debate whether diversification dynamics can be inferred from phylogenies including both extant and extinct species is ongoing (78; 79) because fossils add another axis of information while requiring another rate parameter (the fossilization rate). In our analyses, extinction rate estimates of almost zero events per million years are clearly unrealistic for crocodylians as the fossil record shows strong evidence of species extinctions (37; 32; 33; 35). Furthermore, zero extinction rate estimates can be due to incomplete species sampling at the present (6; 9; 7), as, for example, if true species are missed because of cryptic diversity. Thus, the apparent but slight slowdown of speciation rates and zero extinction rate estimates of the species-level phylogeny are likely an artefact and a signal of underestimated present-day diversity.

Based on our results of our extant-level diversification analyses, future studies in Crocodylia focusing on diversification dynamics including both palaeontological and neontological data should consider cryptic diversity in order to obtain a better understanding of macroevolutionary processes underlying crocodylian evolution. It is worth noting, nevertheless, that the current identification of species in the fossil record has been fundamentally questioned upon the discovery of cryptic species in extant crocodylians, as it demonstrates the need of major reassessment of our understanding concerning crocodylian morphological variation (25). This is furthermore complicated if we consider identifying cryptic diversity in the fossil record, which is unlikely due to inherent sampling bias. Strategies for addressing cryptic diversity in the fossil record need to be further investigated by, for example, computational approaches (i.e. simulation studies) in order to allow a more comprehensive understanding of crocodylian speciation through time. This is a complex albeit fundamental task to be addressed in future macroevolutionary diversification dynamics studies in Crocodylia.

### 4.2 Sampling of extant diversity in macroevolutionary studies

The integration of cryptic diversity in macroevolutionary studies has been advocated for in biodiversity science (80; 81; 13; 8). The impact of cryptic diversity in diversification analysis has been recently discussed in a study focused on freshwater fishes, which has demonstrated that addressing cryptic diversity is critical for model selection in diversification analysis and to the understanding of speciation dynamics (8). A multi-faceted approach for recognizing species (i.e. integrative taxonomy) instead of the sole use of traditional definition of species as morphological diagnosable entities addresses extant diversity in a more comprehensive way as it samples genetically distinct populations that are otherwise overlooked by morphology-only definitions (82; 16; 19; 13). However, as observed in Crocodylia, cryptic lineages detected by molecular studies still lack taxonomic classification (i.e. South American alligatorids (figure 1a,b,c, (52; 53; 20; 19; 55; 21)). Difficulties on taxonomic classifications are due to the complexity of species delimitation under traditional taxonomic practices (e.g.(83)). Those discrepancies in addition to the fact that genetic species discovery methods are recently applied to Crocodylia leads to an inadequate representation of crocodylian extant diversity by macroevolutionary studies (e.g. (56; 5; 30)). As demonstrated in our results, a more comprehensive sampling of extant crocodylian diversity —evidenced by the inclusion of cryptic diversity— affects outcomes of macroevolutionary analyses considerably. We have sampled cryptic crocodylian diversity by using sequences of the mitochondrial gene *ctyb* –a widely used robust gene for phylogenetic inference and delimitation of evolutionary lineages in South American alligatorids (52; 20; 19; 55)–, although investigation of crocodylian diversification dynamics under different underlying genomic data is encouraged.

Studies on cryptic lineages discovery are central for biodiversity estimates, conservation efforts, and taxonomic implications (84; 18; 19; 85). For example, the scientific integration of combining different data sources and approaches to address diversity have a direct impact on taxonomic acts and furthermore in conservation perspectives (80; 24; 18; 20; 86; 19; 85; 87; 88). Many of the detected cryptic lineages are facing several anthropogenic threats including mining, agriculture, and pollution, for instance (19; 54). Habitat loss leads to population fragmentation and reduction, therefore requiring increased efforts on conservation strategies. Many of cryptic crocodylian species are yet not named and globally inserted in a Last Concern status of IUCN Red List due wide distribution of species complexes. While including cryptic diversity to better infer the true diversification dynamics in crocodylians is essential, a future loss of these threatened cryptic species makes the impact on biodiversity loss even greater than without considering these cryptic species. Efforts on molecular studies mapping genetic diversity among populations (e.g. (20; 19; 21; 55; 54)) are critical for the understanding of population structuring and evolutionary aspects, considering that a comprehensive molecular data sampling is a fundamental component for macroevolutionary analyses.

## 5 Conclusions

Our study is the first to address cryptic diversity in a diversification dynamics analysis in Crocodylia. We reported significant increase in speciation and net-diversification rates towards the recent time when using a time-calibrated molecular phylogeny that includes cryptic diversity. These results are in contrast with previously published diversification dynamics studies —that exclude cryptic diversity— in which slowdowns of speciation and net-diversification rates were observed in Crocodylia instead. Therefore, we emphasise the importance of species status and estimates of present-day diversity on diversification dynamics. Furthermore, we raise attention for a multi-faceted approach for addressing extant crocodylian diversity in future studies, specifically for the incorporation of well-established genetic clusters detected by molecular studies. Note that here we estimated diversification rate estimates drawn from molecular phylogenies using a single mitochondrial gene, therefore further investigation of diversification rates under additional or alternative underlying genomic data is encouraged. Additionally, genetic information from cryptic diversity coupled with morphological data and temporal information from the fossil record is yet to be further investigated in future macroevolutionary studies in Crocodylia, as it can contribute to better understand rates of speciation, extinction and diversification, and drivers underlying the evolutionary history of Crocodylia (and more inclusive clades). However, how to appropriately sample the fossil record in respect to cryptic diversity remains an open question. Our results represent an alternative interpretation of diversification rates in Crocodylia and reinforces the recognition of cryptic diversity for a more comprehensive discussion on crocodylian evolution.

## Supporting information

electronic supplementary material

## 7 Acknowledgments

We thank xx and yy for comments that helped improve the manuscript. We thank the members of the Höhna Lab at LMU Munich for helpful discussions.

