## Supplementary material for "The effects of cryptic diversity on diversification dynamics analyses in Crocodylia": electronic supplementary material

January 22, 2025

**DOI:**

### 1 Extended methods: fossil calibration set 2

We used two sets of primary calibrations based on the crocodylian fossil record and extensive literature review to estimate a time-calibrated phylogenies of Crocodylia. The first calibration set, here referred as calibration set 1, follows the same calibrations used by Darlim and Höhna (2024). Calibration set 1 represents a selection of well-justified fossils, and it relies strictly on the best practices of Parham et al. (2012) especially concerning phylogenetic consensus of extinct taxa (i.e. confident phylogenetic affinities based on a review of morphological, molecular and total evidence phylogenetic inferences in Crocodylia). Therefore, calibration set 1 is a more conservative approach for assigning fossil to the selected nodes when compared to calibration set 2 (see below). A full justification for all fossils used for the inference of a time-calibrated tree using the calibration set 1 can be found in Darlim and Höhna (2024). The calibration set 1 includes: *Brachychampsa sealeyi* (Crocodylia), *Protocaiman peligrensis* (Alligatoridae), *Centenariosuchus gilmorei* (crown Caimaninae), *Alligator thomsoni* (crown Alligatorinae), *Kentisuchus spenceri* and *Marroscosuchus zennaroi* (Longirostres), and *Euthecodon arambourguii* (Osteolaeminae). A list containing the age range and settings for calibration densities is provided at Table S2.

Conversely, many key extinct taxa in Crocodylia are subject to considerable uncertainty regarding their phylogenetic placement. To accommodate discussions on the putative affinities of taxa concerning the crocodylian nodes herein calibrated, and to obtain a comparative time-calibrated phylogeny for diversification dynamics analyses, we explored how divergence ages are estimated when using an alternative set of calibrations (i.e. calibration set 2). This alternative set of calibrations consists of fossil taxa with contentious phylogenetic affinities concerning crown groups. We reinforce that the purpose of the selection of an additional set of calibrations is to explore and evaluate different divergence age estimates in Crocodylia, and how the different sets of calibrations affect the results of diversification dynamics analyses. Below we discuss the selected fossils calibrations set 2 for the following nodes: Crocodylia (Node 1), Alligatoridae (Node 2), crown Caimaninae (Node 3), and Crocodylidae (Node 6). Node numbers follow Darlim and Höhna (2024).

#### 1.1 Crocodylia (Node 1)

We selected *Portugalosuchus azenhae* from the upper Cenomanian of Portugal as an alternative calibration for the minimum age of Crocodylia. The phylogenetic position of *Portugalosuchus azenhae* is contentious in two main aspects: (i) *Portugalosuchus* is often retrieved with close affinities to 'thoracosaurs' (Lee and Yates, 2018; Mateus et al., 2019; Rio and Mannion, 2021; Darlim et al., 2022; Salas-Gismondi et al., 2022; Puértolas-Pascual et al., 2023; Burke et al., 2024a,b). 'Thoracosaurs' consists of a polyphyletic group that has received considerably attention in recent phylogenetic studies in Crocodylia especially concerning early evolution of gavialoids and the age of Crocodylia. Long-snouted taxa such as *Thoracosaurus* spp., *Eosuchus* spp., *Eothoracosaurus mississippiensis*, *Argochampsa krebsi*, and *Portugalosuchus azenhae* are commonly referred as 'thoracosaurs' and usually recovered with affinities to gavialoids (Lee and Yates, 2018; Rio and

Mannion, 2021; Darlim et al., 2022; Salas-Gismondi et al., 2022; Burke et al., 2024a,b). However, Lee and Yates (2018) demonstrated that under a total-evidence tip-dating approach which incorporates stratigraphy into phylogenetic reconstruction, 'thoracosaurs' are not recovered as part of Crocodylian. The study of Lee and Yates (2018) further discusses that homoplastic characters due to convergence on skull and snout shape for aquatic feeding suggest a close relationship between 'thoracosaurs' and crown gavialoids in undated phylogenetic analyses based on morphology-only, and even combined with molecular information of extant crocodylians in different phylogenetic approaches, such as maximum parsimony and undated Bayesian inference. Thoracosaurs as non-crocodylians was corroborated in subsequent total-evidence tip-dating analyses in Crocodylia (Darlim et al., 2022; Salas-Gismondi et al., 2022). Nevertheless, phylogenetic position of 'thoracosaurs' awaits reassessment (Rio and Mannion, 2021; Mateus et al., 2019; Puértolas-Pascual et al., 2023). (ii) Additionally, the age of *Portugalosuchus azenhae* (ca. 95 mya) is significantly older than the oldest unambiguous extinct crocodylian (i.e. *Brachychamsa sealeyi*, ca. 82 mya). Thus, using *Portugalosuchus azenhae* as a calibration for Crocodylia favors unjustifiable long ghost lineages at the root of the clade (see Darlim et al. (2022) for a detailed discussion). Thus, we selected the Cenomanian age of *Portugalosuchus azenhae* to explore divergence ages in Crocodylia and the impact on diversification dynamics analyses considering the relevance of *Portugalosuchus azenhae* and 'thoracosaurs' on discussing crocodylian evolution.

The holotype of *Portugalosuchus azenhae* comes from Late Cretaceous deposits (Upper Cenomanian) of the Lower member (Unit B) of Tentúgal Formation. This strata has been widely studied in the fields of micropaleontology and invertebrate palaeontology, in which biostratigraphic correlation (i.e. standard Biozone of *Calycoceras naviculare*) suggests an age of about 95 Ma. A detailed explanation concerning the age (and palaeofauna) of the site where *Portugalosuchus azenhae* was found is provided by Mateus et al. (2019). We specified a soft-bound uniform-normal calibration density between 95 - 115 millions years, with a hard lower bound and a 5% probability of being older with standard deviation of 2.5 millions years.

#### 1.2 Alligatoridae (node 2)

We selected *Brachychampsa sealeyi* as the alternative fossil calibration for Alligatoridae. Differing from our analysis using the calibration set 1, in which *Brachychampsa sealeyi* was used to calibrate the age of Crocodylia as recommended in previous studies (Walter et al., 2022; Darlim et al., 2022; Green et al., 2014), in the calibration set 2 we addressed an alternative phylogenetic position commonly retrieved in the literature as discussed below. *Brachychampsa sealeyi* has been commonly retrieved as one of the early divergent lineages in Alligatoroidea (Norell et al., 1994; Brochu, 1999, 2004, 2010, 2011; Hastings et al., 2013; Pinheiro et al., 2013; Cossette and Brochu, 2018; Lee and Yates, 2018; Godoy et al., 2021; Walter et al., 2022). Conversely, *Brachychampsa sealeyi* has also been recovered with close affinities with stem caimanines (Salas-Gismondi et al., 2015; Bona et al., 2018; Cossette and Brochu, 2018; Stocker et al., 2021; Rio and Mannion, 2021; Bona et al., 2024; Cossette and Tarailo, 2024). The disputed phylogenetic position of *Brachychampsa sealyi* within Alligatoroidea, although unambiguous regarding its alligatoroid affinities, illustrates the complex and still unresolved understanding of the evolution of early alligatoroids (Bona et al., 2018; Godoy et al., 2021; Stocker et al., 2021; Walter et al., 2022). Therefore, considering that the phylogenetic position of *Brachychampsa sealyi* is intimately related to the age of Crocodylia and crown alligatorids, here we use the age interval 75-81 mya to calibrate the split between the two main alligatoid lineages, Alligatorinae and Caimaninae.

A depositional age for the locality of *Brachychampsa sealeyi* (Menefee Formation, New Mexico) was estimated between 75-81 mya based on U-Pb dating (Dickinson and Gehrels, 2008) (see Walter et al. (2022) for extensive review). We specified a soft-bounded uniform-normal distribution between 75 and 90 million years, with 5% probability of being older with standard deviation 2.5 million years.

#### 1.3 crown Caimaninae (node 3)

Phylogenetic relationships of *Bottosaurus harlani* within crown Caimaninae is contentious and directly affects the age of the crown group, therefore making *Bottosaurus harlani* a good candidate for our comparative analysis. *Bottosaurus harlani* has been recovered as a stem caimanine (Walter et al., 2022), and also within the crown-Caimaninae, specifically closer related to *Paleosuchus* spp. (Cossette and Brochu, 2018; Massonne et al., 2019; Cidade et al., 2020; Stocker et al., 2021; Rio and Mannion, 2021; Cossette, 2021; Cossette and Tarailo, 2024; Bona et al., 2024). However, the latter hypothesis (i.e. a crown position for *Bottosaurus harlani*) is ambiguously supported (Cossette and Brochu, 2018; Cossette, 2021; Walter et al., 2022). In a extensive review of the phylogenetic relationships of extinct caimanines, Walter et al. (2022) emphasized the homoplastic synapomorphies and elaborated on the lack of key Caimaninae synapomorphies in *Bottosaurus harlani*. Additionally, close affinities of *Bottosaurus harlani* with *Paleosuchus* spp. suggests long ghost lineages and complex biogeography for the origin of crown Caimaninae. *Bottosaurus harlani* specimens are from Late Cretaceous/early Paleocene of North America, whereas oldest unambiguous fossil crown caimaninae comes from the early Miocene of Central America (Walter et al., 2022; Cossette and Brochu, 2018). Therefore, *Bottosaurus harlani* as a crown-Caimaninae suggests a complicated set of dispersal and back dispersal events from North to South America (for a detailed review see Walter et al. (2022); Cossette (2021). The ambiguous phylogenetic support of *Bottosaurus harlani* in Caimaninae illustrates uncertainty regarding the phylogeny of Cretaceous caimanines and poor understanding of morphological evolution among early alligatoroids (Hastings et al., 2013; Walter et al., 2022; Cossette, 2021).

*Bottosaurus* is an example of key crocodylian taxa that directly affect our interpretation of the origin of a crown clade. Therefore, we herein explored how setting the age of *Bottosaurus harlani* as a minimum constraint for crown Caimaninae compares to robust fossil calibrations os calibration set 1 (i.e. *Centenariosuchus gilmorei*). *Bottosaurus harlani* comes from the Hornerstown Formation in North America, dating from of the Upper Cretaceous to Lower Paleocene (ca. 65 – 66 mya, (Miller et al., 2010; Cossette and Brochu, 2018; Cossette, 2021)). We specified a lognormal distribution with an offset of 66.5 millions years and median of 4 millions years with standard deviation of 0.75.

#### 1.4 Crocodylidae (node 6)

In our calibration set 1, we used the age of *Euthecodon arambourgii* and *Brochuchus pigotti* to calibrate Osteolaeminae (i.e. *Osteolaemus tetraspis* and all crocodylians closer to it than to *Crocodylus niloticus*, (Brochu, 2003)). In that case, the split between *Osteolaemus* spp. and *Mecistops* spp. Phylogenetic support of *Euthecodon arambourgii* and *Brochuchus pigotti* composing Osteolaeminae is generally robust (Brochu, 2007; Brochu and Storrs, 2012; Lee and Yates, 2018; Cossette et al., 2020; Rio and Mannion, 2021; Hekkala et al., 2020; Brochu et al., 2022). Conversely, some of those analyses (Cossette et al., 2020; Brochu et al., 2022) have also retrieved unresolved phylogenetic position of *Euthecodon arambourgii* and *Brochuchus pigotti* in relation to *Mecistops cataphractus* and *Crocodylus* spp., although within Crocodylidae. To address potential uncertainties, we decided to use the age of *Euthecodon arambourgii* as the minimum age of Crocodylidae —node that includes the last common ancestor of *Crocodylus niloticus* and *Osteolaemus tetraspis* and all of its descendants (Brochu, 2003)— for the calibration set 2 (Table S2).

As described in Darlim and Höhna (2024), geochronology of *Brochuchus pigotti* locality in the Hiwegi Formation (Kenya) suggest an age of ca. 18 mya (Conrad et al., 2013; Peppe et al., 2011), consistent with the interval of 16–20 mya (Burdigalian, Miocene) suggested from biostratigrapy of the type locality of *Euthecodon arambourgii* (Tchernov and Couvering, 1978; Buffetaut, 1979; Tchernov, 1986; Conrad et al., 2013) We specified a soft-bound uniform-normal distribution between 16–33 million years, with 5% probability to be younger or older with standard deviation of 1.0.

***Table S1**: GenBank accession numbers and length (bp) of the mitochondrial gene Cytb sequences for the crocodylian species used in the analyses of the present study. Haplotypes are marked with an asterisk.*

| Species | Accession number | *Cytb* length (bp) | Source |
|---|---|---|---|
| *Alligator mississippiensis* | EU496863 | 1159 | Venegas-Anaya et al. (2008) |
| *Alligator sinensis* | JF315320 | 1221 | Oaks (2011) |
| *Melanosuchus niger* | JF315312 | 1221 | Oaks (2011) |
| *Caiman latirostris* (Parana)* | MT472907 | 1117 | Roberto et al. (2020) |
| *Caiman latirostris* (Doce)* | MT472890 | 1120 | Roberto et al. (2020) |
| *Caiman latirostris* (São Francisco)* | MT472882 | 1120 | Roberto et al. (2020) |
| *Caiman crocodilus* (Orinoco/Negro)* | MT472945 | 1200 | Roberto et al. (2020) |
| *Caiman crocodilus* (Brazilian Shield)* | MT473047 | 1140 | Roberto et al. (2020) |
| *Caiman crocodilus* (Amazonia)* | MT473213 | 1014 | Roberto et al. (2020) |
| *Caiman crocodilus* (Upper Branco)* | MT473202 | 1161 | Roberto et al. (2020) |
| *Caiman crocodilus* (Southwestern Amazon)* | MT473189 | 1149 | Roberto et al. (2020) |
| *Caiman crocodilus 'fuscus'* | MT473139 | 1149 | Roberto et al. (2020) |
| *Caiman crocodilus cf. 'fuscus'* | EU 496839 | 1150 | Venegas-Anaya et al. (2008) |
| *Caiman crocodilus 'chiapasius'* | EU 496845 | 1150 | Venegas-Anaya et al. (2008) |
| *Caiman yacare* (Pantanal)* | MT472976 | 1168 | Roberto et al. (2020) |
| *Caiman yacare* (Bolivia/Madeira)* | MT473184 | 1149 | Roberto et al. (2020) |
| *Paleosuchus trigonatus* (West Amazon)* | OQ859174 | 1020 | Hernández-Rangel et al. (2024) |
| *Paleosuchus trigonatus* (Southwest Amazon)* | MH757611 | 1020 | Bittencourt et al. (2019) |
| *Paleosuchus trigonatus* (West Orinoco)* | OQ859166 | 1020 | Hernández-Rangel et al. (2024) |
| *Paleosuchus trigonatus* (East Amazon)* | MH757689 | 1020 | Bittencourt et al. (2019) |
| *Paleosuchus trigonatus* (Upper Branco)* | OQ859175 | 1020 | Hernández-Rangel et al. (2024) |
| *Paleosuchus trigonatus* (Coastal)* | MH757470 | 1020 | Bittencourt et al. (2019) |
| *Paleosuchus palpebrosus* (Madeira)* | MH846438 | 1097 | Muniz et al. (2019) |
| *Paleosuchus palpebrosus* (Bolivia)* | MH846476 | 1097 | Muniz et al. (2019) |
| *Paleosuchus palpebrosus* (Orinoco)* | OQ859159 | 1089 | Hernández-Rangel et al. (2024) |
| *Paleosuchus palpebrosus* (Amazonas)* | OQ859165 | 1089 | Hernández-Rangel et al. (2024) |
| *Paleosuchus palpebrosus* (Pantanal)* | MH846500 | 1097 | Muniz et al. (2019) |
| *Crocodylus siamensis* | JF315292 | 1200 | Oaks (2011) |
| *Crocodylus palustris* | JF315283 | 1200 | Oaks (2011) |
| *Crocodylus porosus* | JF315290 | 1200 | Oaks (2011) |
| *Crocodylus novaeguineae* | JF315286 | 1200 | Oaks (2011) |
| *Crocodylus mindorensis* | JF315247 | 1200 | Oaks (2011) |
| *Crocodylus johnstoni* | JF315260 | 1200 | Oaks (2011) |
| *Crocodylus rhombifer* | JF315261 | 1200 | Oaks (2011) |
| *Crocodylus moreletii* | JF315256 | 1200 | Oaks (2011) |
| *Crocodylus intermedius* | JF315259 | 1200 | Oaks (2011) |
| *Crocodylus acutus* | JF315258 | 1200 | Oaks (2011) |
| *Crocodylus suchus** | MT727030 | 1189 | Hekkala et al. (2020) |
| *Crocodylus niloticus* | JF315257 | 1200 | Oaks (2011) |
| *Osteolaemus tetraspis* | JF315287 | 1200 | Oaks (2011) |
| *Osteolaemus osborni** | MN885917 | 1200 | Meredith - direct entry |
| *Mecistops cataphractus* | JF315265 | 1200 | Oaks (2011) |
| *Mecistops leptorhynchus** | MN885915 | 1200 | Meredith - direct entry |
| *Tomistoma schlegelii* | JF315305 | 1197 | Oaks (2011) |
| *Gavialis gangeticus* | JF315302 | 1197 | Oaks (2011) |

***Table S2:*** *List of fossil calibrations for the two calibration sets used for inferring time calibrated topology of Crocodylia. Asterisk on node names in calibration set 2 indicates that alternative fossil calibration used compared to calibration set 1.*

| Node | Calibration set 1 (Darlim and Höhna, 2024) | | | |
|---|---|---|---|---|
| | **Taxon** | **Age (mya)** | **Calibration density** | **Standard deviation (sd)** |
| Node 1, Crocodylia | *Brachychampsa sealeyi* | 75-81 | soft-bound uniform-normal (75-90 mya) | hard lower bound and 5% probability of being older with sd of 2.5 |
| Node 2, Alligatoridae | *Protocaiman peligrensis/Necrosuchus ionensis* | 63.5-65.7 | normal with mean of 67.5 mya | 1.775 |
| Node 3, crown Caimaninae | *Centenariosuchus gilmorei* | 18.06 | lognormal with an offset of 18.06 mya and median of 4 mya | 0.75 |
| Node 4, crown Alligatorinae | *Alligator thomsoni* | 13.6-16.3 | soft-bound uniform-normal (13.6-25 mya) | 5% probability to be younger or older with sd of 1.0 |
| Node 5, Longirostres | *Kentisuchus spenceri/ Maroccosuchus zennaroi* | 48.6 | soft-bound uniform-normal (48.6 - 66 Mya) | 5% probability to be younger or older with sd of 1.0 |
| Node 6, Osteolaeminae | *Euthecodon arambourgii* | 16 | soft-bound uniform-normal (16 - 33 mya) | 5% probability to be younger or older with sd of 1.0 |
| | **Calibration set 2** | | | |
| | **Taxon** | **Age (mya)** | **Calibration density** | **Standard deviation (sd)** |
| Node 1, Crocodylia* | *Portugalosuchus azenhae* | 95 | soft-bound uniform-normal (95-113 mya) | hard lower bound and 5% probability of being older with sd of 2.5 mya |
| Node 2, Alligatoridae* | *Brachychampsa sealeyi* | 75-81 | soft-bound uniform-normal (75-90 mya) | hard lower bound and 5% probability of being older with sd of 2.5 mya |
| Node 3, crown Caimaninae* | *Bottosaurus harlani* | 65-66 | lognormal with an offset of 66.5 mya and median of 4 mya | 0.75 |
| Node 4, crown Alligatorinae | *Alligator thomsoni* | 13.6-16.3 | soft-bound uniform-normal (13.6-25 mya) | 5% probability to be younger or older with sd of 1.0 |
| Node 5, Longirostres | *Kentisuchus spenceri/ Maroccosuchus zennaroi* | 48.6 | soft-bound uniform-normal (48.6 - 66 mya) | 5% probability to be younger or older with sd of 1.0 |
| Node 6, Crocodylidae* | *Euthecodon arambourgii* | 16 | soft-bound uniform-normal (16 - 33 mya) | 5% probability to be younger or older with sd of 1.0 |

### 2 Results

Our time-calibrated phylogenies using fossil calibrations of the two different sets differ quite significantly (Table S3 and Table S4). As expected, divergence age estimates using the calibration set 2 (older calibrations) are considerably older. Older ages are specifically noticeable for Caimaninae in both analyses including and excluding cryptic diversity (mean ages of 69.43 mya and 69.11 mya, respectively), as well as for Alligatoridae (mean ages of 79.96 mya and 84.97 mya). Similarly, divergence age estimates for Crocodylia when using calibration set 2 are older with respect to calibration set 1, extending back to approximately 115 mya. Such age estimate further differs from previous molecular studies (i.e. Oaks (2011); Pan et al. (2021); Darlim and Höhna (2024)). Conversely, the age of Osteolaeminae and Crocodylidae are overall similar in all analyses excluding and including cryptic diversity, and using calibration sets 1 and 2. Interestingly, the results of our diversification dynamics analyses using the species-level and the cryptic-species-level phylogenies for both calibration sets are furthermore overall similar, except for a slight reduction of the 95% highest posterior intervals in the rate estimates when using calibration set 2. As discussed in Walter et al. (2022) and Parham et al. (2012), the use of fossils of uncertain phylogenetic placement contributed to an overestimation of crown ages. As explained in the fossil calibrations section, fossils assigned to the calibration set 2 in our study represent species whose phylogenetic relationships are still debated and await further assessment. Nevertheless, because previous studies have still recovered crown affinities for those fossils, it was suitable for our analysis to explore divergence age estimates upon different calibrations in order to investigate potential effects on diversification dynamics in Crocodylia. It should be noted that, based on the overestimation of node ages recovered by our analyses when using fossil calibration set 2, we recommend the use of fossil in calibration set 1 instead, as they represent more reasonable and justified selections. Nonetheless, it remains crucial to reassess fossil calibrations in future studies aiming to time-calibrate the crocodylian tree, considering new fossil descriptions and advances in crocodylian systematics.

#### 2.1 Phylogenetic inference and diversification analyses using calibration set 1

##### 2.1.1 Convergence assessment of the phylogenetic inference

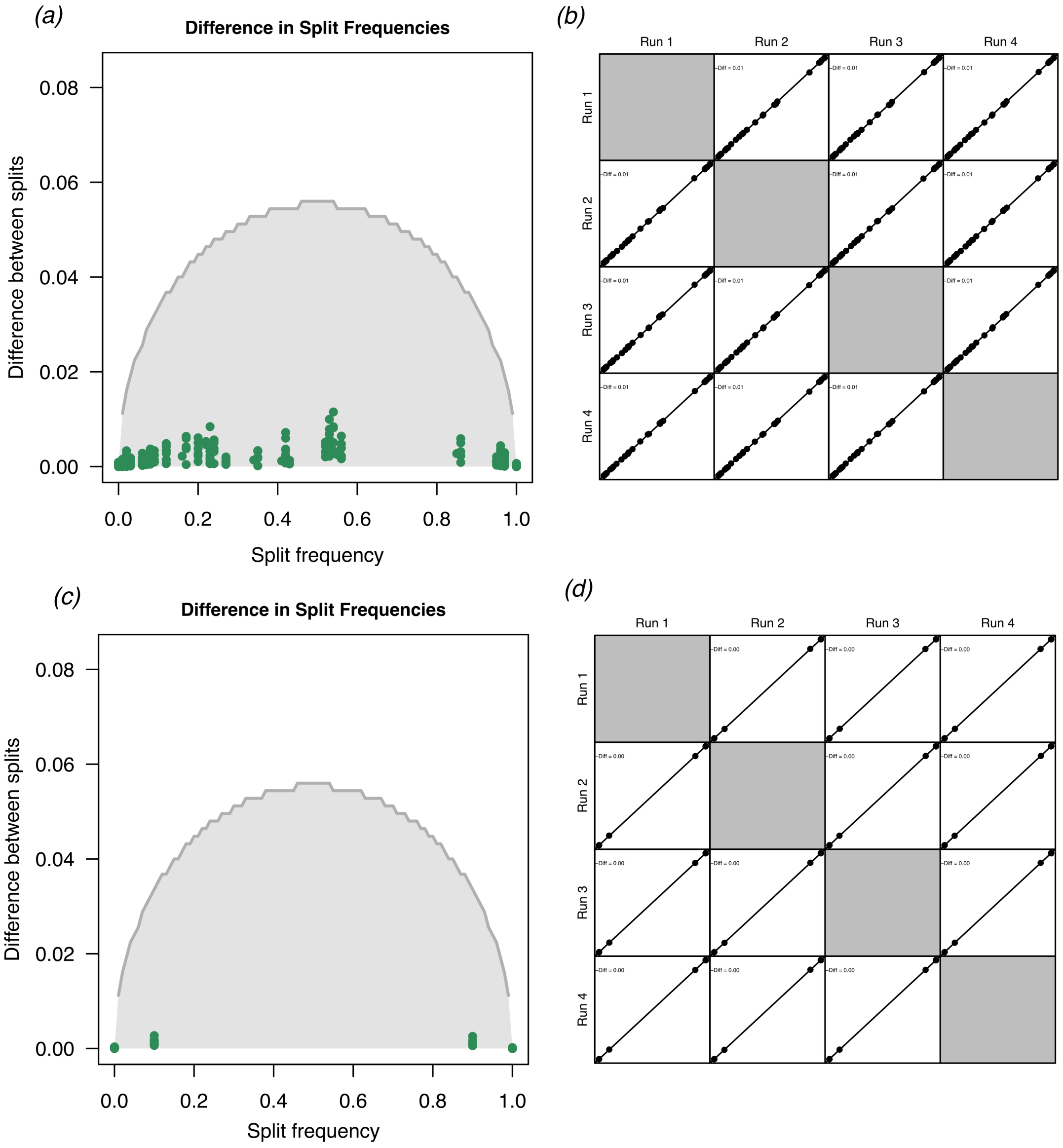

***Figure S1***: *Convergence assessment of the phylogenetic inferences including (a,b) and excluding (c,d) cryptic species using the R package convenience (Fabreti and Höhna, 2022). Split differences between pairwise replicates (a,c). Cummulative split difference compared to expectation when assuming at least 625 independent tree samples (b,d). The results show that our phylogenetic tree estimation converged well, i.e., all 4 replicate MCMC runs converged to the same posterior distribution of trees and splits.*

##### 2.1.2 Divergence age estimates

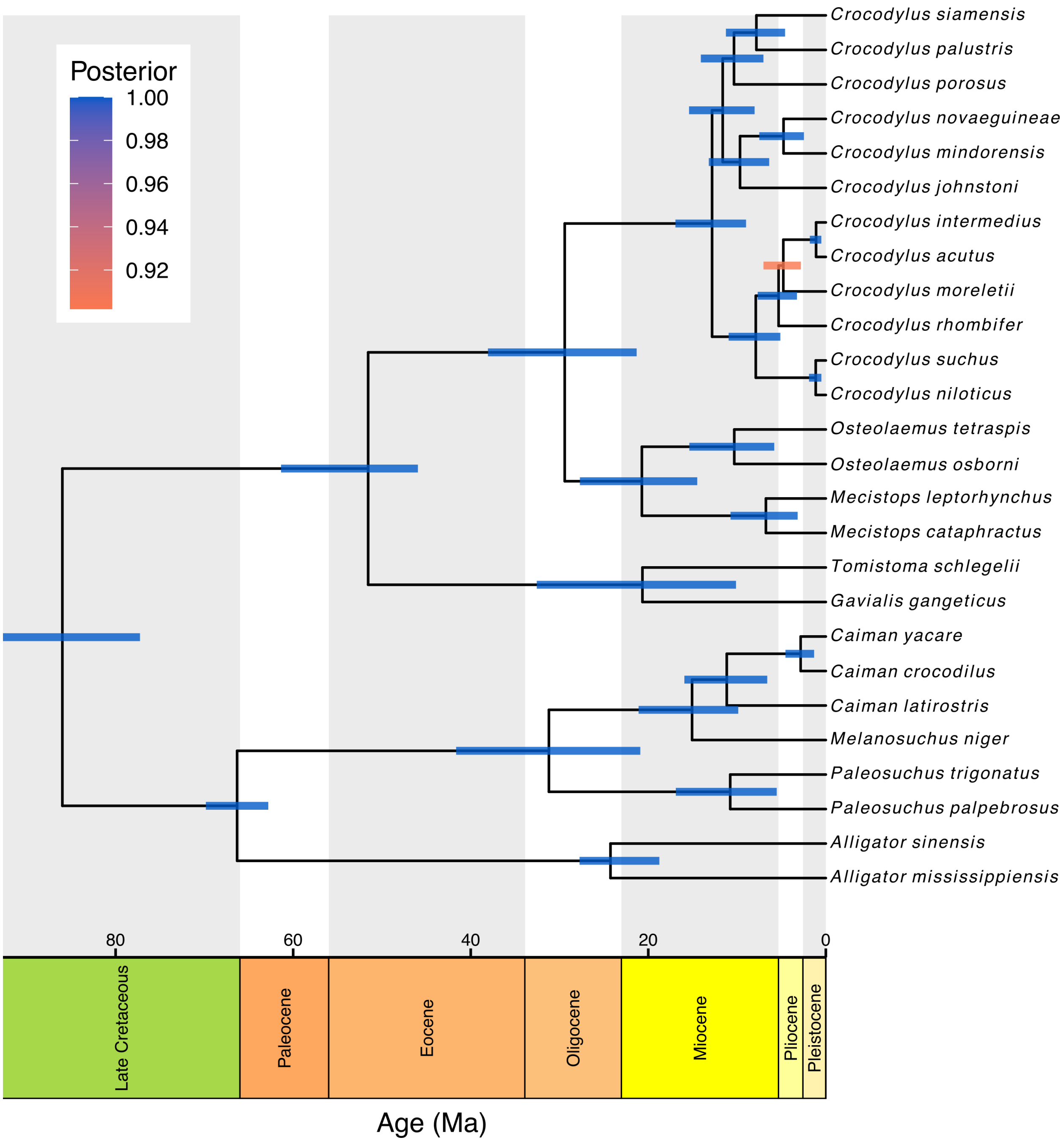

***Figure S2***: *Time-calibrated phylogeny of the species-level tree of Crocodylia using calibration set 1. Node bars indicate 95% highest posterior density interval (HPD) for age estimates. Node bar colors are in accordance to the posterior support index illustrated at the up left corner of the figure. The phylogeny was plotted using the R package RevGadgets (Tribble et al., 2022).*

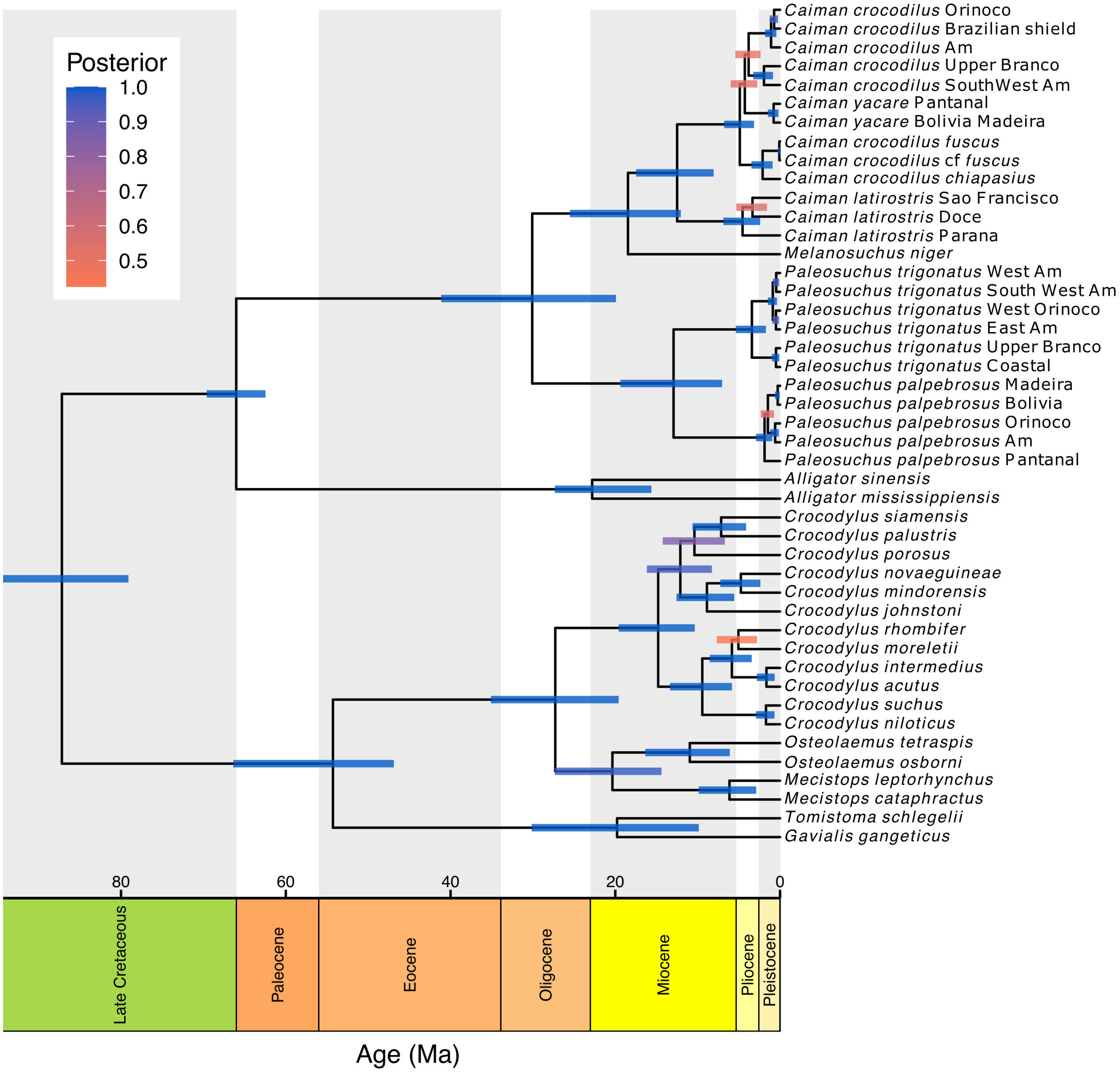

***Figure S3**: Time-calibrated phylogeny of the cryptic-species-level tree of Crocodylia. Node bars indicate 95% highest posterior density interval (HPD) for age estimates. Node bar colors are in accordance to the posterior support index illustrated at the up left corner of the figure. The phylogeny was plotted using the R package RevGadgets (Tribble et al., 2022).*

***Table S3.*** *Node ages estimates of our time-calibrated crocodylian tree using calibration set 1. The table shows a comparison between divergence age estimates of common nodes of analyses including (present study) and excluding cryptic diversity. Node ages are represented in million years before the present (mya) as both means and 95% highest posterior density intervals (HPD). Node numbers are in accordance with those of figure 2 of the main manuscript.*

| Node | MRCA | Cryptic diversity included | | Cryptic diversity excluded | |
|---|---|---|---|---|---|
| | | Node ages | | Node ages | |
| | | Mean | 95% HPD | Mean | 95% HPD |
| 1 | **Crocodylia** | 87.18 | 79.1 - 94.31 | 86 | 77.25 – 92.69 |
| 2 | **Alligatoridae** | 66.01 | 62.46 - 69.6 | 66.31 | 62.8 – 69.83 |
| 3 | **crown-Alligatorinae** | 22.81 | 15.62 - 27.34 | 24.26 | 18.75 - 27.74 |
| 4 | **crown-Caimaninae** | 30.09 | 19.93 - 41.13 | 31.19 | 20.9 – 41.64 |
| 5 | **Jacarea** (*Caiman* spp., *Paleosuchus* spp., *Melanosuchus niger*) | 18.46 | 12.05 - 25.5 | 15.07 | 9.86 – 21.09 |
| 6 | *Caiman* spp. | 12.5 | 8.06 - 17.47 | 11.16 | 6.6 – 15.93 |
| 7 | ***Caiman latirostris* species complex** | 4.55 | 2.42 - 6.89 | - | - |
| 8 | *Ca. latirostris* Doce, *Ca. latirostris* São Francisco | 3.35 | 1.59 - 5.31 | - | - |
| 9 | ***Caiman crocodilus*, *Caiman yacare* species complex** | 4.89 | 3.16 - 6.78 | - | - |
| 10 | *Ca. crocodilus fuscus*, *Ca. crocodilus chiapasius* | 2.12 | 0.9 - 3.47 | - | - |
| 11 | *Ca. crocodilus fuscus*, *Ca. crocodilus* cf. *fuscus* | 0.09 | 0.0 - 0.28 | - | - |
| 12 | *Ca. yacare*, *Ca. crocodilus* | 4.28 | 2.75 - 6 | - | - |
| 13 | ***Caiman yacare* species complex** | 0.78 | 0.2 - 1.48 | - | - |
| 14 | *Ca. crocodilus* Brazilian shield, *Ca. crocodilus* Orinoco, *Ca. crocodilus* Amazonia, *Ca. crocodilus* SouthWest Amazon *Ca. crocodilus* Upper Branco | 3.81 | 2.37 - 5.43 | - | - |
| 15 | *Ca. crocodilus* SouthWest Amazon, *Ca. crocodilus* Upper Branco | 1.95 | 0.86 - 3.25 | - | - |
| 16 | *Ca. crocodilus* Brazilian shield, *Ca. crocodilus* Orinoco, *Ca. crocodilus* Amazonia | 1.09 | 0.45 - 1.82 | - | - |
| 17 | *Ca. crocodilus* Brazilian shield, *Ca. crocodilus* Orinoco | 0.72 | 0.25 - 1.27 | - | - |
| 18 | *Paleosuchus* spp. | 12.93 | 7.03 - 19.39 | 10.77 | 5.53 – 16.9 |
| 19 | ***Paleosuchus trigonatus* species complex** | 3.42 | 1.72 - 5.37 | - | - |
| 20 | *Pa. trigonatus* Coastal, *Pa. trigonatus* Upper Branco | 0.49 | 0.1 - 0.98 | - | - |
| 21 | *Pa. trigonatus* East Amazon, *Pa. trigonatus* West Orinoco, *Pa. trigonatus* SouthWest Amazon, *Pa. trigonatus* West Amazon | 0.88 | 0.37 - 1.48 | - | - |
| 22 | *Pa. trigonatus* SouthWest Amazon, *Pa. trigonatus* West Amazon | 0.47 | 0.11 - 0.89 | - | - |
| 23 | *Pa. trigonatus* East Amazon, *Pa. trigonatus* West Orinoco | 0.5 | 0.12 - 0.94 | - | - |
| 24 | ***Paleosuchus palpebrosus* species complex** | 1.88 | 0.95 - 2.93 | - | - |
| 25 | *Pa. palpebrosus* Amazonas, *Pa. palpebrosus* Orinoco *Pa. palpebrosus* Bolivia, *Pa. palpebrosus* Madeira | 1.48 | 0.74 - 2.34 | - | - |
| 26 | *Pa. palpebrosus* Amazonas, *Pa. palpebrosus* Orinoco | 0.6 | 0.15 - 1.16 | - | - |
| 27 | *Pa. palpebrosus* Bolivia, *Pa. palpebrosus* Madeira | 0.29 | 0.04 - 0.61 | - | - |
| 28 | **Longirostres** | 54.28 | 46.88 - 66.38 | 51.55 | 45.95 – 61.36 |
| 29 | **Crocodylidae** | 27.28 | 19.59 - 35.1 | 29.41 | 21.31 – 38.06 |
| 30 | **Osteolaeminae** (*Osteolaemus* spp., *Mecistops* spp.) | 20.35 | 14.39 - 27.34 | 20.72 | 14.48 – 27.7 |
| 31 | *Osteolaemus tetraspis*, *Osteolaemus osborni* | 10.96 | 6.1 - 16.35 | 10.31 | 5.81 - 15.38 |
| 32 | *Mecistops cataphractus*, *Mecistops leptorhynchus* | 6.14 | 2.89 - 9.86 | 6.75 | 3.17 - 10.74 |
| 33 | **Crocodylinae** (*Crocodylus* spp.) | 14.8 | 10.35 - 19.6 | 12.81 | 8.99 – 16.94 |
| 34 | Australasian *Crocodylus* spp. (*Cr. porosus*, *Cr. siamensis*, *Cr. palustris*, *Cr. johnstoni*, *Cr. novaeguinae*, *Cr. mindorensis*) | 12.11 | 8.27 - 16.16 | 11.62 | 8.02 – 15.41 |
| 35 | *Cr. porosus*, *Cr. siamensis*, *Cr. palustris* | 10.39 | 6.7 - 14.25 | 10.34 | 7.03 – 14.08 |
| 36 | *Cr. siamensis*, *Cr. palustris* | 7.15 | 4.12 - 10.63 | 7.82 | 4.59 – 11.25 |
| 37 | *Cr. johnstoni*, *Cr. novaeguinae*, *Cr. mindorensis* | 8.88 | 5.57 - 12.57 | 9.66 | 6.38 – 13.19 |
| 38 | *Cr. novaeguinae*, *Cr. mindorensis* | 4.76 | 2.4 - 7.26 | 4.77 | 2.47 – 7.5 |
| 39 | African + Neotropical *Crocodylus* spp. (*Cr. acutus*, *Cr. intermedius*, *Cr. moreletii*, *Cr. rhombifer*, *Cr. niloticus*, *Cr. suchus*) | 9.44 | 5.85 - 13.36 | 7.89 | 5.12 – 10.95 |
| 40 | *Cr. acutus*, *Cr. intermedius*, *Cr. moreletii*, *Cr. rhombifer* | 5.85 | 3.44 - 8.54 | 5.32 | 3.25 – 7.67 |
| 41 | *Cr. moreletii*, *Cr. rhombifer* | 5.06 | 2.79 - 7.68 | - | - |
| 42 | *Cr. acutus*, *Cr. intermedius* | 1.65 | 0.65 - 2.83 | 1.1 | 0.51 – 1.8 |
| 43 | *Cr. niloticus*, *Cr. suchus* | 1.71 | 0.68 - 2.93 | 1.14 | 0.5 - 1.9 |
| 44 | **Gavialidae** (*Gavialis gangeticus*, *Tomistoma schlegelii*) | 19.78 | 9.86 - 30.15 | 20.66 | 10.12 – 32.56 |

##### 2.1.3 Diversification dynamics analysis

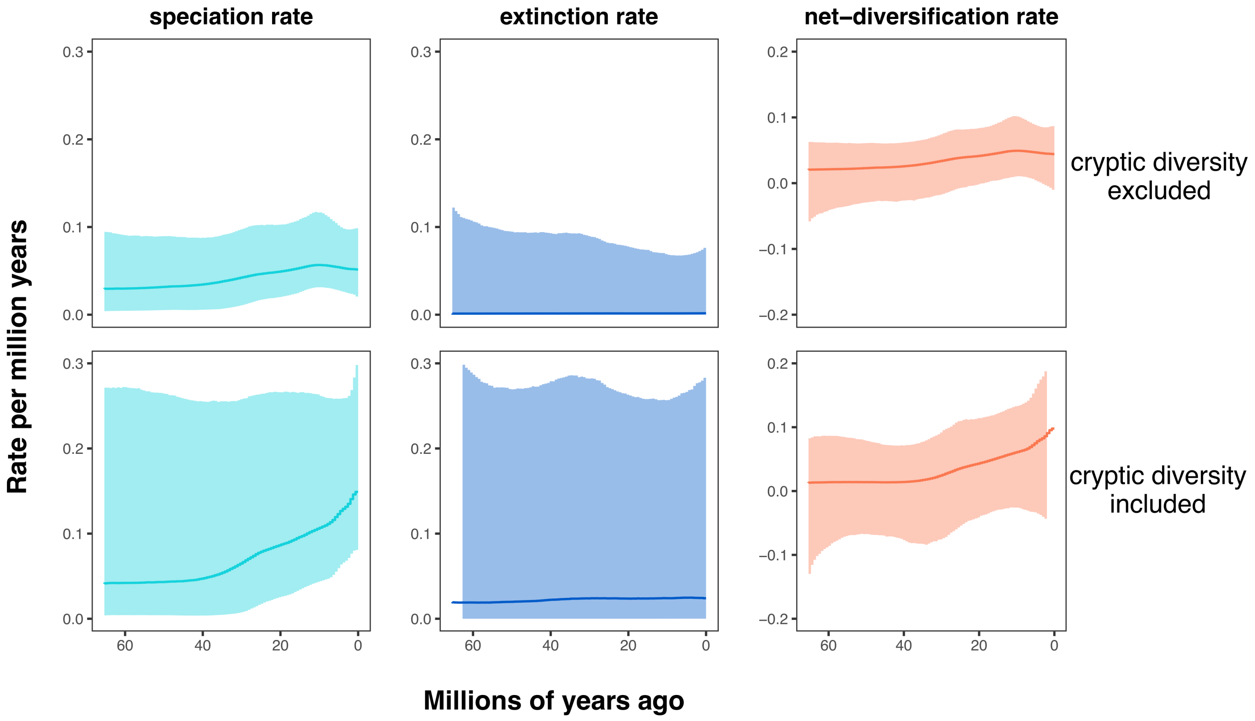

***Figure S4**: Diversification dynamics analyses in Crocodylia showing estimated speciation, extinction, and net-diversification rates across time based on data sets excluding (top row) and including (bottom row) cryptic diversity under calibration set 1. Analyses were performed under the piecewise-constant birth-death process with 100 equal-sized intervals between 66 million years ago and the present implemented in* `RevBayes` *(Höhna et al., 2016). Results are summarized and plotted with the R package* `RevGadgets` *(Tribble et al., 2022). Posterior probability of a speciation rate decrease for the species-level analysis between the peak at ~ 10 mya and the present was* $P(\lambda(t=10) > \lambda(t=0)) = 0.61$*, which gives a Bayes factor of* $BF = \frac{P(\lambda(t=10)>\lambda(t=0))}{P(\lambda(t=10)\leq\lambda(t=0))} = 1.56$*, signifying weak support. Posterior probability of a speciation rate increase for the cryptic-species-level analysis between* ~ 40 *mya and the present was* $P(\lambda(t=40) < \lambda(t=0)) = 0.89$*, which gives a Bayes factor of* $BF = \frac{P(\lambda(t=40)<\lambda(t=0))}{P(\lambda(t=40)\geq\lambda(t=0))} = 8.09$*, signifying substantial support. Solid line line represents the median.*

#### 2.2 Phylogenetic inference and diversification analyses using calibration set 2

##### 2.2.1 Convergence assessment of the phylogenetic inference

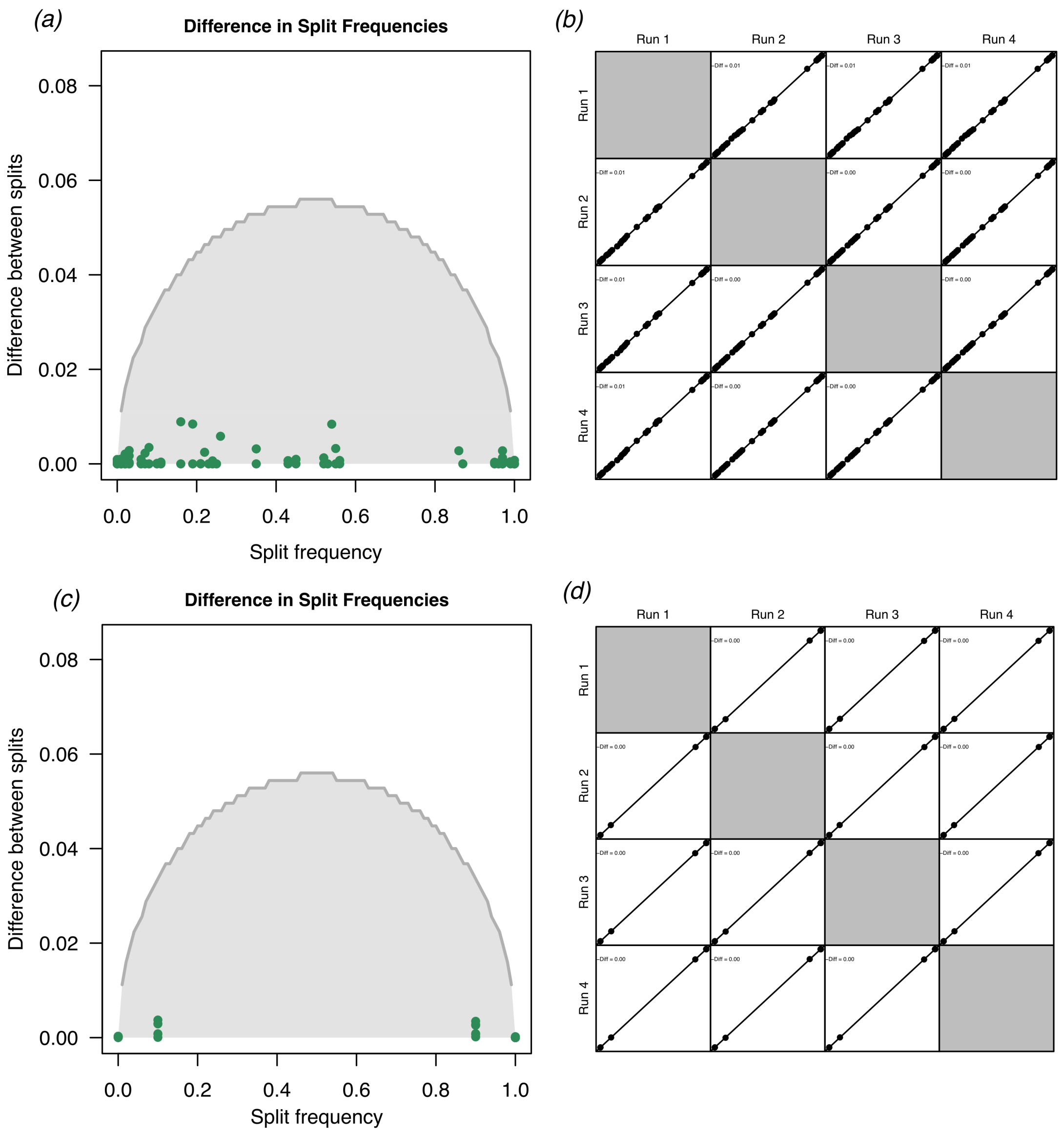

***Figure S5***: *Convergence assessment of the phylogenetic inferences including (a,b) and excluding (c,d) cryptic species using the R package convenience (Fabreti and Höhna, 2022). Split differences between pairwise replicates (a,c). Cummulative split difference compared to expectation when assuming at least 625 independent tree samples (b,d). The results show that our phylogenetic tree estimation converged well, i.e., all 4 replicate MCMC runs converged to the same posterior distribution of trees and splits.*

##### 2.2.2 Divergence age estimates

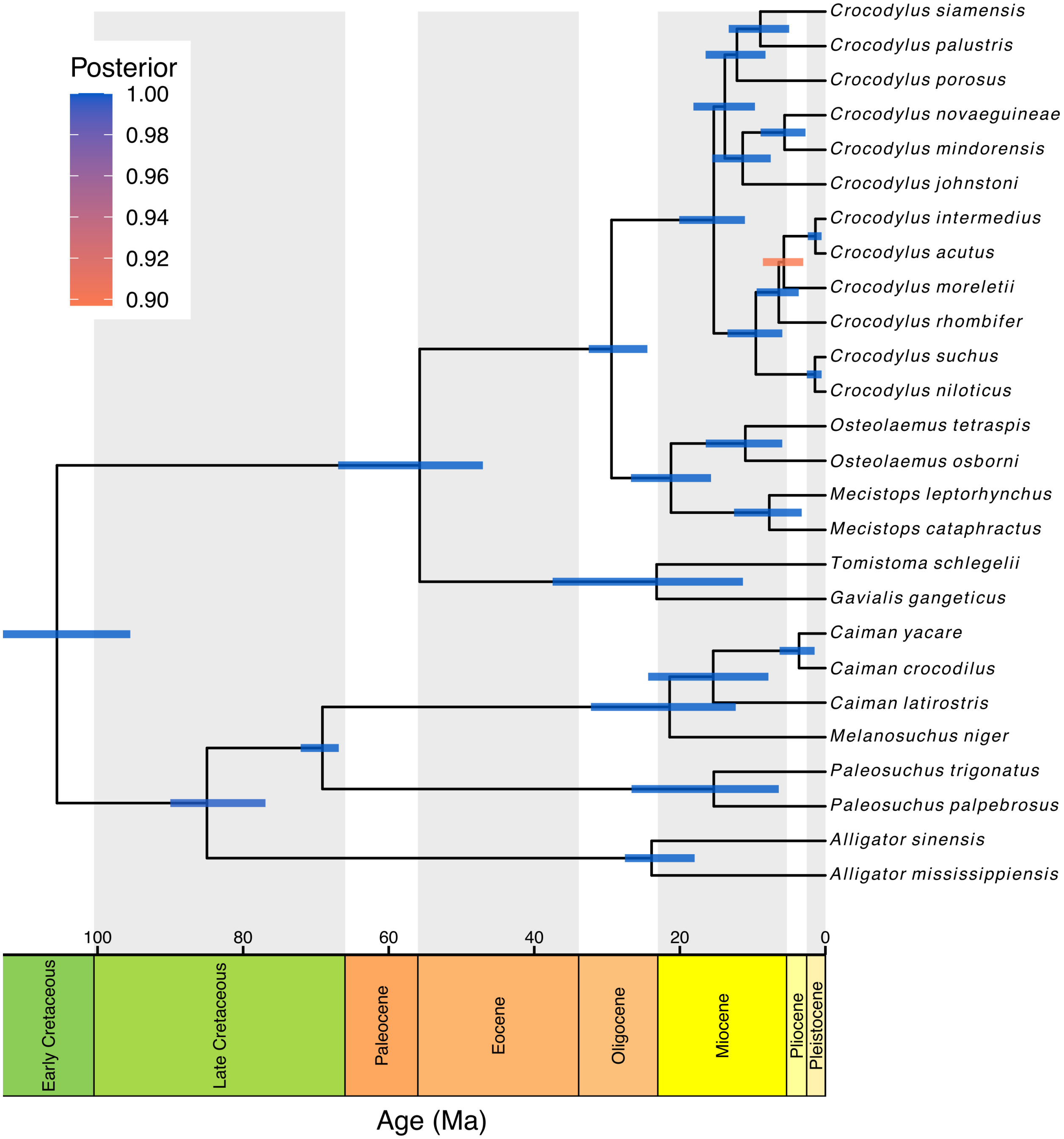

***Figure S6***: *Time-calibrated phylogeny of the species-level tree of Crocodylia using calibration set 2. Node bars indicate 95% highest posterior density interval (HPD) for age estimates. Node bar colors are in accordance to the posterior support index illustrated at the up left corner of the figure. The phylogeny was plotted using the R package RevGadgets (Tribble et al., 2022).*

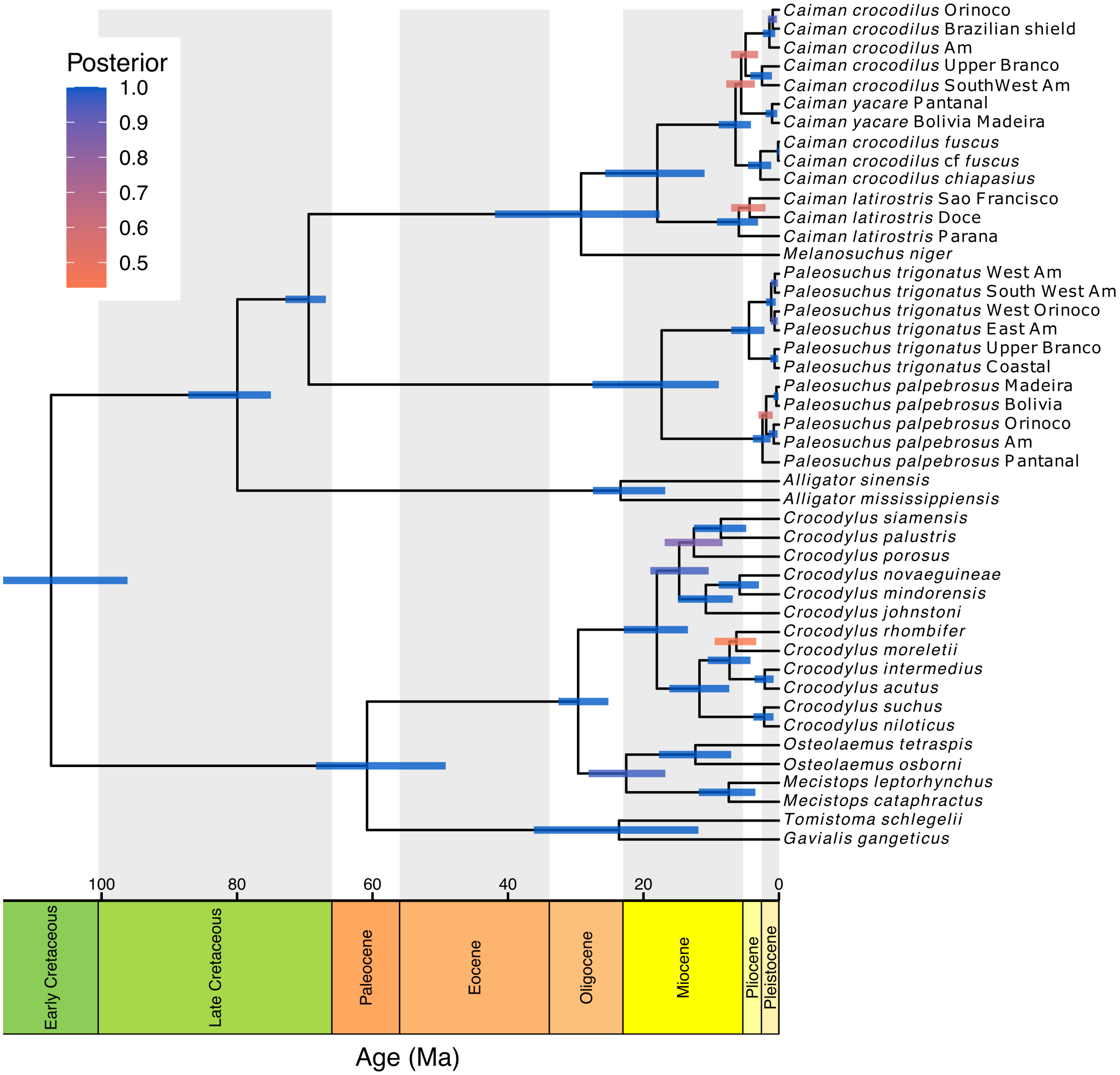

***Figure S7***: *Time-calibrated phylogeny of the cryptic-species-level tree of Crocodylia using calibration set 2. Node bars indicate 95% highest posterior density interval (HPD) for age estimates. Node bar colors are in accordance to the posterior support index illustrated at the up left corner of the figure. The phylogeny was plotted using the R package RevGadgets (Tribble et al., 2022).*

***Table S4.****Node ages estimates of our time-calibrated crocodylian tree using calibration set 2. The table shows a comparison between divergence age estimates of common nodes of analyses including (present study) and excluding cryptic diversity. Node ages are represented in million years before the present (mya) as both means and 95% highest posterior density intervals (HPD).*

| Node | MRCA | Cryptic diversity included | | Cryptic diversity excluded | |
|---|---|---|---|---|---|
| | | Node ages | | Node ages | |
| | | Mean | 95% HPD | Mean | 95% HPD |
| 1 | **Crocodylia** | 107.46 | 96.17 - 114.54 | 105.58 | 95.5 – 113 |
| 2 | **Alligatoridae** | 79.96 | 75 - 87.2 | 84.97 | 76.93 – 90 |
| 3 | **crown-Alligatorinae** | 23.38 | 16.79 - 27.46 | 23.88 | 17.97 - 27.55 |
| 4 | **crown-Caimaninae** | 69.43 | 66.89 - 72.86 | 69.11 | 66.86 – 72.09 |
| 5 | **Jacarea**<br>(*Caiman* spp., *Paleosuchus* spp., *Melanosuchus niger*) | 29.21 | 17.61 - 41.92 | 21.4 | 12.32 – 32.21 |
| 6 | *Caiman* spp. | 17.95 | 10.97 - 25.65 | 15.42 | 7.83 – 24.36 |
| 7 | ***Caiman latirostris* species complex** | 5.92 | 3.08 - 9.16 | - | - |
| 8 | *Ca. latirostris* Doce, *Ca. latirostris* São Francisco | 4.32 | 1.98 - 7.06 | - | - |
| 9 | ***Caiman crocodilus*, *Caiman yacare* species complex** | 6.41 | 4.13 - 8.89 | - | - |
| 10 | *Ca. crocodilus fuscus*, *Ca. crocodilus chiapasius* | 2.73 | 1.14 - 4.57 | - | - |
| 11 | *Ca. crocodilus fuscus*, *Ca. crocodilus* cf. *fuscus* | 0.11 | 0.0 - 0.36 | - | - |
| 12 | *Ca. yacare*, *Ca. crocodilus* | 5.57 | 3.55 - 7.78 | - | - |
| 13 | ***Caiman yacare* species complex** | 1.01 | 0.26 - 1.96 | - | - |
| 14 | *Ca. crocodilus* Brazilian shield, *Ca. crocodilus* Orinoco,<br>*Ca. crocodilus* Amazonia, *Ca. crocodilus* SouthWest Amazon<br>*Ca. crocodilus* Upper Branco | 4.92 | 3.07 - 7.04 | - | - |
| 15 | *Ca. crocodilus* SouthWest Amazon, *Ca. crocodilus* Upper Branco | 2.5 | 1.03 - 4.22 | - | - |
| 16 | *Ca. crocodilus* Brazilian shield, *Ca. crocodilus* Orinoco,<br>*Ca. crocodilus* Amazonia | 1.41 | 0.55 - 2.36 | - | - |
| 17 | *Ca. crocodilus* Brazilian shield, *Ca. crocodilus* Orinoco | 0.92 | 0.3 - 1.63 | - | - |
| 18 | *Paleosuchus* spp. | 17.32 | 8.87 - 27.53 | 15.34 | 6.41 – 26.64 |
| 19 | ***Paleosuchus trigonatus* species complex** | 4.42 | 2.14 - 7.04 | - | - |
| 20 | *Pa. trigonatus* Coastal, *Pa. trigonatus* Upper Branco | 0.64 | 0.12 - 1.28 | - | - |
| 21 | *Pa. trigonatus* East Amazon, *Pa. trigonatus* West Orinoco,<br>*Pa. trigonatus* SouthWest Amazon, *Pa. trigonatus* West Amazon | 1.14 | 0.47 - 1.9 | - | - |
| 22 | *Pa. trigonatus* SouthWest Amazon, *Pa. trigonatus* West Amazon | 0.6 | 0.15 - 1.15 | - | - |
| 23 | *Pa. trigonatus* East Amazon, *Pa. trigonatus* West Orinoco | 0.64 | 0.16 - 1.21 | - | - |
| 24 | ***Paleosuchus palpebrosus* species complex** | 2.42 | 1.22 - 3.85 | - | - |
| 25 | *Pa. palpebrosus* Amazonas, *Pa. palpebrosus* Orinoco<br>*Pa. palpebrosus* Bolivia, *Pa. palpebrosus* Madeira | 1.88 | 0.91 - 3.02 | - | - |
| 26 | *Pa. palpebrosus* Amazonas, *Pa. palpebrosus* Orinoco | 0.77 | 0.18 - 1.5 | - | - |
| 27 | *Pa. palpebrosus* Bolivia, *Pa. palpebrosus* Madeira | 0.37 | 0.05 - 0.8 | - | - |
| 28 | **Longirostres** | 60.81 | 49.19 - 68.34 | 55.75 | 47.06 – 66.96 |
| 29 | **Crocodylidae** | 29.64 | 25.18 - 32.55 | 29.39 | 24.46 – 32.52 |
| 30 | **Osteolaeminae**<br>(*Osteolaemus* spp., *Mecistops* spp.) | 22.54 | 16.77 - 28.11 | 21.21 | 15.72 – 26.72 |
| 31 | *Osteolaemus tetraspis*, *Osteolaemus osborni* | 12.33 | 7.05 - 17.71 | 11 | 5.93 - 16.44 |
| 32 | *Mecistops cataphractus*, *Mecistops leptorhynchus* | 7.4 | 3.47 - 11.83 | 7.71 | 3.26 - 12.56 |
| 33 | **Crocodylinae**<br>(*Crocodylus* spp.) | 18.02 | 13.44 - 22.9 | 15.34 | 11.06 – 20.09 |
| 34 | Australasian *Crocodylus* spp.<br>(*Cr. porosus*, *Cr. siamensis*, *Cr. palustris*,<br>*Cr. johnstoni*, *Cr. novaeguinae*, *Cr. mindorensis*) | 14.72 | 10.37 - 18.99 | 13.82 | 9.68 – 18.13 |
| 35 | *Cr. porosus*, *Cr. siamensis*, *Cr. palustris* | 12.56 | 8.32 - 16.87 | 12.15 | 8.23 – 16.46 |
| 36 | *Cr. siamensis*, *Cr. palustris* | 8.57 | 4.83 - 12.54 | 8.94 | 4.98 – 13.29 |
| 37 | *Cr. johnstoni*, *Cr. novaeguinae*, *Cr. mindorensis* | 10.79 | 6.82 - 14.92 | 11.36 | 7.51 – 15.56 |
| 38 | *Cr. novaeguinae*, *Cr. mindorensis* | 5.8 | 2.95 - 8.88 | 5.61 | 2.73 – 8.91 |
| 39 | African + Neotropical *Crocodylus* spp.<br>(*Cr. acutus*, *Cr. intermedius*, *Cr. moreletii*,<br>*Cr. rhombifer*, *Cr. niloticus*, *Cr. suchus*) | 11.73 | 7.33 - 16.18 | 9.56 | 5.92 – 13.46 |
| 40 | *Cr. acutus*, *Cr. intermedius*, *Cr. moreletii*, *Cr. rhombifer* | 7.29 | 4.19 - 10.51 | 6.41 | 3.67 – 9.45 |
| 41 | *Cr. moreletii*, *Cr. rhombifer* | 6.28 | 3.35 - 9.5 | - | - |
| 42 | *Cr. acutus*, *Cr. intermedius* | 2.1 | 0.78 - 3.6 | 1.37 | 0.53 – 2.42 |
| 43 | *Cr. niloticus*, *Cr. suchus* | 2.17 | 0.8 - 3.76 | 1.44 | 0.52 - 2.55 |
| 44 | **Gavialidae**<br>(*Gavialis gangeticus*, *Tomistoma schlegelii*) | 23.61 | 11.89 - 36.17 | 23.19 | 11.33 – 37.48 |

##### 2.2.3 Diversification dynamics analysis

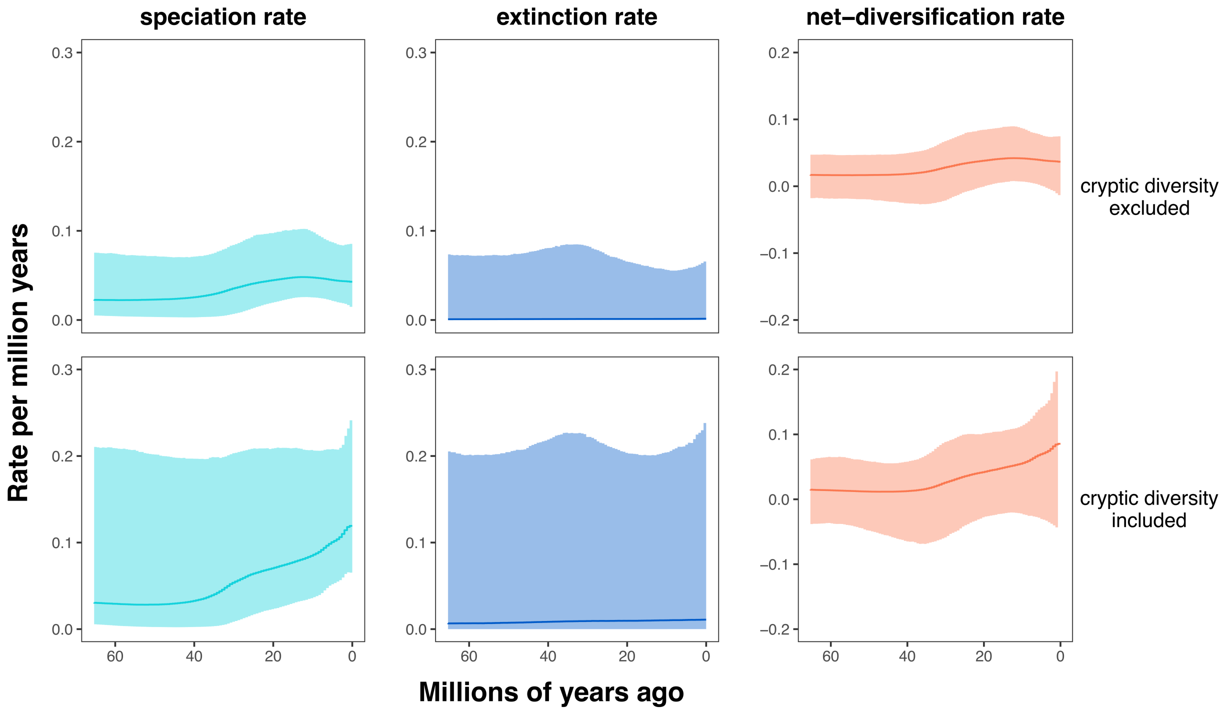

***Figure S8**: Diversification dynamics analyses in Crocodylia showing estimated speciation, extinction, and net-diversification rates across time based on data sets excluding (top row) and including (bottom row) cryptic diversity under calibration set 2. Analyses were performed under the piecewise-constant birth-death process with 100 equal-sized intervals between 66 million years ago and the present implemented in* `RevBayes` *(Höhna et al., 2016). Results are summarized and plotted with the R package* `RevGadgets` *(Tribble et al., 2022). Posterior probability of a speciation rate decrease for the species-level analysis between the peak at* $\sim 12.5$ *mya and the present was* $P(\lambda(t = 12.5) > \lambda(t = 0)) = 0.63$*, which gives a Bayes factor of* $BF = \frac{P(\lambda(t=12.5)>\lambda(t=0))}{P(\lambda(t=12.5)\leq\lambda(t=0))} = 1.70$*, signifying weak support. Posterior probability of a speciation rate increase for the cryptic-species-level analysis between* $\sim 40$ *mya and the present was* $P(\lambda(t = 40) < \lambda(t = 0)) = 0.89$*, which gives a Bayes factor of* $BF = \frac{P(\lambda(t=40)<\lambda(t=0))}{P(\lambda(t=40)\geq\lambda(t=0))} = 8.09$*, signifying substantial support. Solid line line represents the median.*
